# T Cell Repertoire Evolution After Allogeneic Bone Marrow Transplantation: An Organizational Perspective

**DOI:** 10.1101/326744

**Authors:** Jeremy A. Meier, Mohamed Fawaz, Hamdi Abdeen, Jason Reed, Christopher G. Kanakry, Leo Luznik, Amir A. Toor

**Author notes:** Correspondence: Amir A. Toor, MD. Professor of Medicine, 1300 East Marshall Ave, Richmond, VA, 23298.

## Abstract

High throughput sequencing (HTS) of human T cell receptors has revealed a high level of complexity in the T cell repertoire. In an attempt to correlate T cell reconstitution with clinical outcomes several measures of T cell repertoire complexity have emerged. However, the associations identified are of a broadly statistical nature, not allowing precise modeling of outcomes based on T cell repertoire development in clinical contexts such as following bone marrow transplantation (BMT). Previous work demonstrated that there is an inherent, mathematically definable order observed in the T cell population that is conserved in a diverse group of donors, and which is perturbed in recipients following BMT. Herein, we use a public database of human leukocyte antigen matched related-donor and recipient T cell receptor (TCR) β sequences to further develop this methodology. TCR β sequencing from unsorted T cells and sorted T cell subsets isolated from peripheral blood samples from BMT donors and recipients show remarkable conservation and symmetry of VJ segment usage in the clonal frequencies, linked to the organization of the gene segments along the TCR locus. This TCR β VJ segment translational symmetry is preserved post-transplant, and even in cases of acute GVHD (aGVHD), suggesting that GVHD occurrence represents a polyclonal donor T cell response to recipient antiges. We also observe that the complexity of the repertoire is significantly diminished after BMT and is not restored even years out post-transplant. The results here provide a new method of quantifying and characterizing post-transplant T cell repertoire reconstitution by further analyzing the mathematical rules governing TCR usage in the context of BMT. This approach may allow for a new means to correlate clinical outcomes with the evolving T cell repertoire post-transplant.

## Introduction

Bone marrow transplantation (BMT) from a human leukocyte antigen (HLA)-matched or HLA haploidentical donor, provides a potentially curative modality in disorders of hematopoiesis as well as hematological malignancies. ^1^ Despite substantial advances in HLA matching and transplant procedures, mortality and morbidity following BMT remain unacceptable, in large part due to graft versus host disease (GVHD) stemming from alloreactivity and variable immune reconstitution contributing to infections. ^2^ This means that donor recipient pairs with equivalent level of histocompatibility have disparate outcomes with associated differences in T cell reconstitution.

T cell reconstitution following BMT is a crucial factor in determining clinical outcomes. Though progress has been made in better understanding this process, ^3^ the complexities inherent to immune recovery after BMT are still elusive. This is particularly the case for the T cell pool, which not only mediates GVHD, but is also crucial to the graft versus leukemia effect (GvL) of an allograft. High throughput sequencing has emerged as an invaluable tool in monitoring the T cell population, ^4,5^ and has provided insight into how the T cell repertoire might evolve following BMT. ^6,7,8^ The T cell repertoire is comprised of millions of T cell clones, each bearing unique T cell receptors (TCR) made up of distinct α and β subunits. These unique TCR subunits are generated by the recombination of gene segments on α and β T cell receptor loci, termed TRA and TRB, and located on chromosomes 14q and 7q respectively. ^9,10^ Each of the TCR loci has variable (V), joining (J) and constant (C) gene segments, and in the case of TRB, there are diversity (D) segments as well. One each of the V and J, and if present, D gene segments on these loci are recombined during T cell ontogeny to yield unique VDJ rearranged, T cell receptor a and b subunits (Supplementary Figure 1). The unique receptor subunits together confer an incredible diversity to the human T cell repertoire enabling recognition of millions of potential antigens, including minor histocompatibility antigens, presented on either matched or mismatched HLA molecules in the context of allogeneic BMT. Recognition of antigens by TCRs on T cells leads to corresponding, antigen specific T cell clonal expansion driving an immune response, which depending on the circumstances may or may not lead to the desired outcomes.

In the case of GVHD (GVHD), high-throughput sequencing has been utilized to identify and track alloreactive T cells.^11,12^ Interpretation of the resulting data has been challenging given not only the unique antigens present in each individual capable of driving this process, but also because of a diverse pool of potentially pathogenic T cell clones. However some trends have emerged; for example, acute GVHD has been associated with a restricted clonal diversity in the T cell population. ^7,13,14^ This has been evident when assessing tissue infiltrating T cells in cases of aGVHD,^15,16^ which often are distinct from patient to patient, and may or may not correlate with the T cells in circulation.^8,16,17^ However, other studies have suggested that acute GVHD may lead to a more diverse T cell phenotype, ^6^ ultimately suggestive of study and transplant type dependency of the observations. Aside from GVHD, other factors, such as cytomegalovirus (CMV) reactivation or serostatus, have also been associated with immune recovery and T cell receptor clonality post-transplant. ^8,18^

As novel treatment strategies emerge to limit post-transplant side effects such as acute or chronic GVHD and maximize the therapeutic potential of allografting, a deeper quantitative understanding of the biology of T cell reconstitution is necessary. This may be possible by interpreting TCR high-throughput sequencing data in terms of the VDJ recombination process. Previous work has demonstrated that lymphocyte recovery post-transplant follows the rules of population growth as in a dynamical system.^19–21^ In such a system, antigen-driven T cell clones grow in proportion to the antigen affinity of the relevant TCR, accounting for environmental constraints such as cytokine effect. This may be used to predict adverse outcomes and simulate alloreactivity using information from the antigen driven T cell clonal array. Thus examining the T cell repertoire after BMT may give valuable insights into GVHD pathophysiology. When evaluating the organization of the repertoire in terms of T cell clones containing various TRB V and J segments, it was noted that the T cell repertoire was self-similar, hence fractal in nature in BMT donors, and tended to be restored to this state with time after BMT suggestive of an inherent order to determining TCR usage.^22^ This order may in fact derive from the organization of the gene segments along the DNA molecules, ^9,23^ which would allow for some level of predictability in terms of T cell reconstitution.

In this paper, using a publicly available database of TCR sequences from BMT donors and recipients, ^8^ the composition of the T cell repertoire and its evolution over time after transplant is further characterized. A novel method of comprehensively analyzing the immune system through high-throughput sequencing is presented. The mathematical properties of the T cell repertoire as these relate to the quantitative aspects of TRB recombination process are analyzed. The impact that maladaptive processes such as acute GVHD may have on the underlying T cell repertoire is also measured.

## Materials and Methods

### Patients and Sequencing Data

Patient sequencing data in this study was a subset from those included in the original IRB-approved multi-institutional prospective clinical trial (NCT00809276) as previously described. ^8,24^ Sequencing results were accessed through the publicly available repository through Adaptive Biotechnologies (Seattle, WA) (https://clients.adaptivebiotech.com/pub/Kanakry-2016-JCIInsight. We included all sixteen patients for which there were sequencing data on both donor (n=16) and corresponding recipient at 1 month (n=9), 2-3 months (n=9), 1 year (n=15), and/or 3+ years (n=3) posttransplant. Per the original study design^24^ patients underwent a myeloablative-conditioning regimen consisting of once daily IV busulfan and fludarabine. The subset of patients included in this analysis all underwent HLA matched-related allogeneic BMT and received post-transplant cyclophosphamide (days +3 and +4) as single-agent GVHD prophylaxis. Acute GVHD was scored as previously described^24^ using Modified Keystone Criteria.

### Samples and TCR Sequencing

Genomic DNA from peripheral blood samples was obtained as previously described in the original study^8^ with sequencing of *TRB* loci at survey level resolution through Adaptive Biotechnologies using the immunoSEQ Platform. Data were reported as copy number for specific clones. The immunoSEQ Platform combines multiplex PCR with high-throughput sequencing and a sophisticated bioinformatics pipeline (including data normalization) for analysis.^25,26^ For T-cell subsets (CD3^+^CD4^+^ and CD3^+^CD8^+^), fluorescence assisted cell sorting (BD FACSAria II) was performed prior to external sequencing as described.^8^ TRB sequencing analyses of those previously published and publicly accessible data are reported in this paper.

### Self-similarity and Symmetry analysis

Self-similarity is a mathematical property of many natural systems, where the structure, or geometry of an object appears similar at different levels of magnification, following the same organizational rules. Common examples of this are the appearance of a tree or the pulmonary airways, where branching patterns are sustained over several orders of division from the trunk to the leaves in the former, and the trachea to the alveoli in the latter. These relationships are quantitatively described by logarithmic scaling and are characterized by a fractal dimension which remains similar no matter what level of magnification the structure is evaluated at, within the limits of that natural system. For visualizing the self-similarity in the T cell repertoire at different levels of organization, such as the frequency of unique TRB-J & -VJ segment containing T cell clones, relative proportional distribution (RPD) graphs were generated as previously described.^22^ Briefly, total counts for unique VJ recombinations were analyzed with V and J segments organized according to their position along the genome at the TCR β locus relative to the 5′ centromeric end of the TRB gene.^9^ Pseudogenes were included in the analysis bringing the total number of possible V segments to sixty-five with the potential to recombine with any of the thirteen J segments found at the TCR β locus. Data were visualized using Microsoft Excel and represent all TCR β recombinations for that particular sample. Each ring in the graph consists of a different J segment (J1.1→J2.7) broken up by the proportion to which each V segment is recombined with each J segment for a total of 100% usage. For clarity of understanding, in this paper we use the term clonal frequency for TCR genomic DNA copy number.

Self-similar structures are usually symmetric. Symmetry refers to the property that an object appears the same when viewed from different perspectives, or it retains its properties after different operations have been performed on it. For example, a circle appears similar when rotated through different angles, and an equilateral triangle similarly appears the same when rotated by 120°, the laws of physics are invariant across time and space. Human anatomy demonstrates several example of reflection symmetry, with the morphology of structures such as the extremities, the brain and lungs demonstrating this symmetry, and at an embryological level, even structures such as the heart and the GI tract develop from symmetrical precursor organs. T cell receptor recombination process yields VJ recombined subunits which have a similar frequency of specific J segments recombining with V segments, with the recombination frequencies maintained between individuals^9^. This may be considered as an example of translational symmetry, where when viewed from the frame of reference of a unique J segment (or vice versa, of unique V segment) the recombination frequencies with each of the V segments (or vice versa, with each J segment) across the TRB locus would always remain the same (Figure 1). To determine this, radar plots were generated by taking the clonal frequency of individual VJ recombination events plotted on the logarithmic scale. Each line in the radar plot represents a distinct J segment with unique V segment recombinations plotted as the graph progresses across the TRB locus. The corresponding rings in the radar plots denote changing magnitude as it progresses.

**Figure 1:**
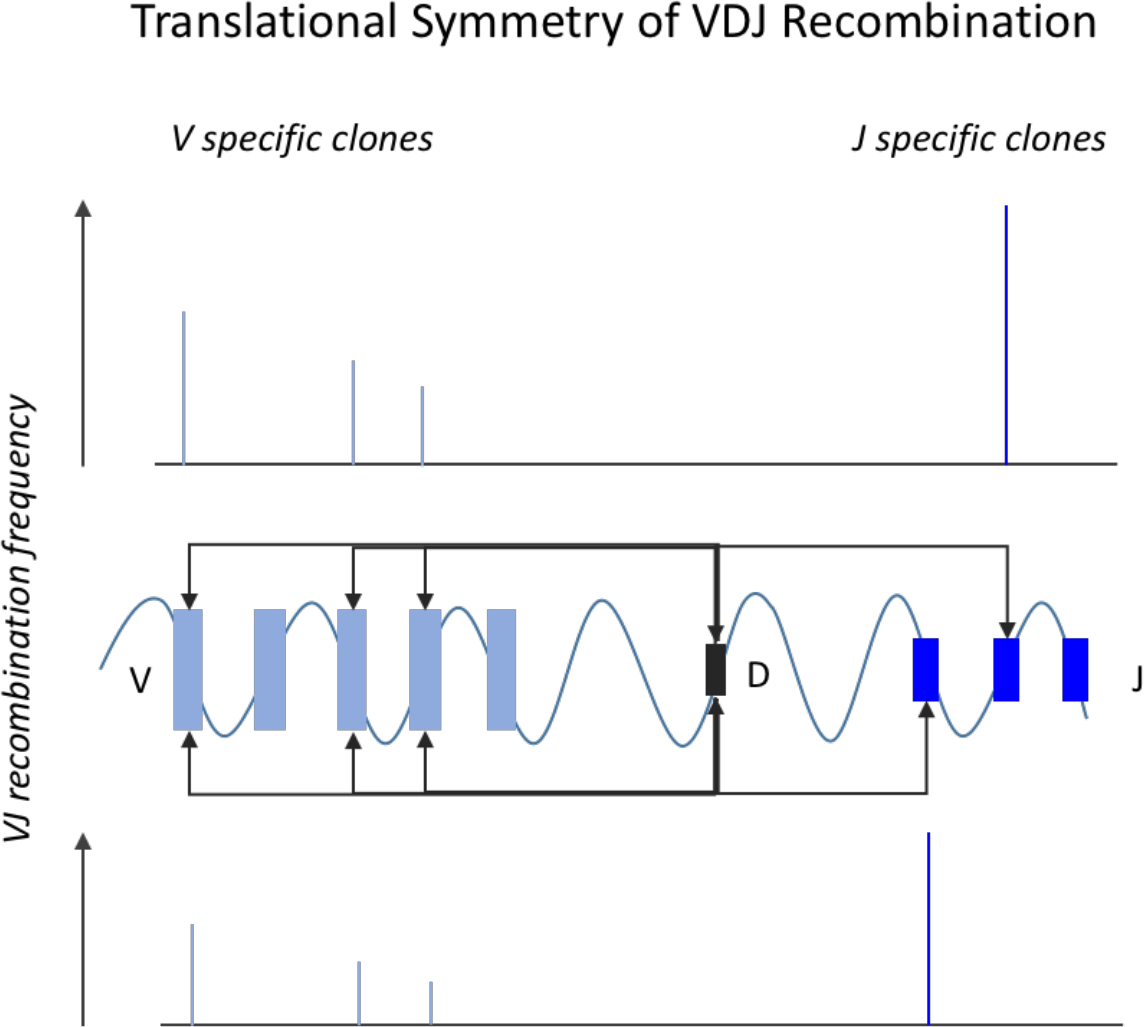
Translational symmetry of TRB VDJ recombination. J segments recombine with specific V segments in similar proportions and vice versa. Bar graphs depict the clonal frequency of specific rearrangements involving the V segments, which vary in absolute magnitude but retain relative proportions when recombining with specific J segments, with D segment acting as an intermediate step.

### Recombination Potentials of TRB V & J segments

The recombination potential is a method of quantifying the translational symmetry of the TRB J and V segments while accounting for complexity of the T cell repertoire as a whole. A challenge in accounting for symmetry in the representation of TRB gene segments is the periodic nature of the gene loci across a very long stretch of DNA. In order to calculate this, the effect of scaling bias (genomic distance between gene segments) has to be eliminated. This may be accomplished by accounting for the helical nature of DNA molecules and calculating the angular coordinates of each TRB segment as previously described.^9^ Briefly, DNA is considered as a propagating wave with each turn of the DNA helix corresponding with 2π radians in terms of angular distance across the TRB locus. Given that there are 10.4 nucleotides per turn of DNA,^27^ the angular coordinates of successive TCR gene segments can be determined by the following equation, where θV is the angular distance of the *k*^*th*^ TRB V (or J segment) from TRB D2 segments and *N* represents the total number of segments

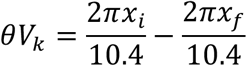

where *X*_*i*_ represents the 5’ initial nucleotide of the TRB-D2 segment and *X*_*f*_ represents the 3’ final nucleotide of the respective V or J segments. The angular distance to the D segments was used in this instance because of the sequence of TRB recombination, which follows a J → D and DJ → V sequence.

The recombination potential of each gene segment can then be derived by taking the average of the recombination potential of each J or V segment between donors and recipients,

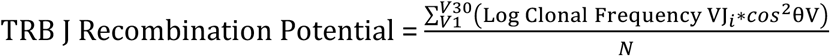

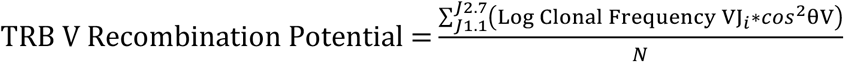

The *cos*^2^θV periodic function eliminates the bias introduced by the genomic position of TRB gene segment on the locus and taking the logarithm of the clonal frequency scales the variability of clonal frequency across the gene segments being studied. N is the total number of gene segments (V or J). These transformations allow an unbiased view of the contribution of each locus to the repertoire complexity. These analyses were stratified based on acute GVHD or recipient CMV serostatus where indicated.

### Pearson Correlation Coefficient

TCR sequence rearrangements (based on productive frequency) between donor and recipient or intra-recipient comparisons were compared using a pairwise scatter plot. Linear regression analysis was performed using immunoSEQ ANALYZER (Adaptive Technologies) on common TRB rearrangements within the two pools with R-values used for analysis. Donor and recipient comparisons were segregated based on aGVHD history or recipient CMV serostatus. Corresponding heat maps were generated based on the Pearson Correlation coefficient comparing donor to recipient or intra-recipient TCR sequences at indicated times post-transplant on the range of −1 to 1.

### Log Rank Plots

Log-log plots of assigned rank versus relative clonal frequency of VJ clones was performed as previously described.^22^ Briefly, the relative clonal frequency was calculated by taking the count total of each unique TCR β VJ recombination and dividing by the sum of all sequence counts for that particular pool to determine the relative usage of that clone. Assigned ranks were based on the percent frequency (*f*) of that clone in the population according to the following schema:

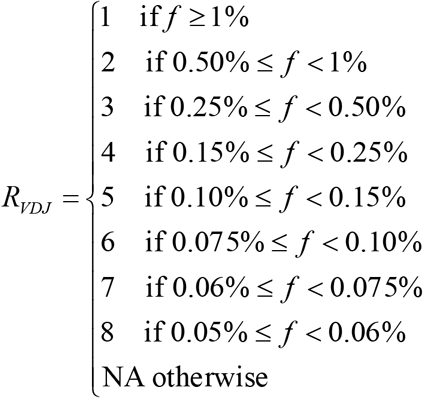

The absolute value of the slope (*a*) of the resulting linear regression lines (*y*=*a* log *x* + log *k*) was reflective of self-similarity of the TCR repertoire and is considered equivalent to the fractal dimension.

### Clonal Tracking

Individual TRB rearrangements for BMT donors or BMT recipients at 1 month, 2-3 months, or 1-year post-transplant were arranged according to clonal frequency and sorted based on the donor repertoire to identify those clones that made up the top donor ranks (i.e. top 200 most frequent donor clones). The percentage with which those rearrangements were found in the corresponding recipient rank (i.e. top 200 recipient clones) was determined for all times post-transplant. Conversely, those clones that populated the top recipient ranks post-transplant were analyzed for their respective position in the donor pool of clones. The top 5 recipient clones that were also identified in the donor samples were used for this frequency analysis. These included T cell clones observed in the recipient, and ranged in order from clone number 1 to 68 in the recipient, depending on donor-recipient pair. Clonal tracking results were stratified based on aGVHD history or recipient CMV serostatus.

### Clonality, Gini coefficient, and Diversity Index

Clonality and the Gini coefficient for certain samples were determined using the LymphoSeq package of software tools available for download at https://bioconductor.org (Coffey D (2017). *LymphoSeq: Analyze high-throughput sequencing of T and B cell receptors.* R package version 1.6.0) as previously described^8^. These measures are expressed on a scale from 0 to 1 with 1 representing a purely monoclonal population. The inverse of Simpson’s Diversity Index was used to calculate the diversity of TCR sequencing samples as previously described ^6^. It is represented by the following equation, 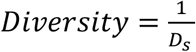 where 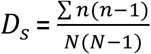 where *n* represents individual unique VJ recombination counts and *N* represents the sum of all VJ recombination counts in that sample.

### Recombination Matrices and Heat Maps

V-J recombination matrices were generated by taking the percent change of total, unique sequence counts for each VJ recombination event when comparing donor to recipient at indicated times post-transplant or by tracking intra-recipient changes over time after transplant. The resulting percent change values were plotted as a heat map with VJ recombination events that were lower in frequency represented in green, and increasing VJ recombinations depicted in red. Scale dependent on particular donor-recipient pair or intra-recipient comparison.

### Statistics

Statistics were performed in GraphPad Prism (Graphpad Software, version 7) using either multiple t-tests (with significance determined using the Holm-Sidak method with alpha=0.05) or a student’s t-test (two-sided, alpha=0.05)

## Results

### Evolution of the T-cell Repertoire after Bone Marrow Transplantation

Using TCR β sequencing data from a group of BMT donor and recipient pairs^8^ (Table 1 & Supplementary Table 1), the self-similar distribution of the T cell repertoire evolution over time was evaluated. Previous work had demonstrated self-similarity across the TCR repertoire amongst BMT donors, and that the T cell repertoire was organized in a fractal order when viewed from the standpoint of TRB gene segment usage.^22^ TCR β VJ recombination clonal frequency (copy numbers) from each donor and recipient time points were arranged in a *n* × *m* matrix in the same order that V and J segments are arranged along the TCR β genomic locus. Clonal frequency of the unique gene segment containing T cell clones consistently demonstrated conservation of the V segment representation across all J segments when the relative proportional distribution (RPD) graphs within the donor pool were examined (Figure 2A, top panel, column 1). Viewing the repertoire in this way gives a means to comprehensively view all TCR recombination frequencies simultaneously. The self-similarity of VJ recombination process within the T cell population is evident by the degree to which each ring in the graph aligns with the others producing a repetitive pattern of TCR β J recombinations with each respective V segment. It is important to note that while not identical, the appearance of these rings is similar across the J segment recombinations visualized. The architecture observed in the donors appears to be distorted when visualized in the recipients early post-transplant, but subsequently recovers with time to assume a more donor-like organization 1 year out from transplantation (Figure 2A, top panel, columns 2-4).

**Table 1:** Summary Patient Characteristics

|  | No GVHD<br>(n=8) | GVHD<br>(n=8) | p value |
| --- | --- | --- | --- |
| Age, median (range) | 47 (22-60) | 59 (38-63) | 0.09962 |
| Sex, female, n (%) | 6 (75%) | 4 (50%) | 0.30302 |
| Disease, n (%) |  |  |  |
| AML | 6 (75%) | 7 (88%) | 0.52218 |
| ALL | 2 (25%) | 0 (0%) | 0.13104 |
| MDS | 0 (0%) | 1 (12%) | 0.30302 |
| Donor Age, median<br>(range) | 49 (17-57) | 57 (35-67) | 0.15559 |
| Donor Sex, female, n(%) | 5 (63%) | 6 (75%) | 0.58920 |
| GVHD |  |  |  |
| aGVHD, n(%) |  | 8 (100%) |  |
| I, n(%) |  | 1(12.5%) |  |
| II, n(%) |  | 6(75%) |  |
| III/IV, n(%) |  | 1(12.5%) |  |
| cGVHD, n(%) |  | 1(12.5%) |  |
| CMV Serology, n(%) |  |  |  |
| D-R<sup>-</sup> | 3 (37.5%) | 2 (25%) | 0.58920 |
| D-R<sup>+</sup> | 1 (12.5%) | 4 (50%) | 0.10524 |
| D<sup>+</sup>R<sup>-</sup> | 2 (25%) | 1 (12.5%) | 0.52218 |
| D<sup>+</sup>R<sup>+</sup> | 2 (25%) | 1 (12.5%) | 0.52218 |
| CMV Reactivation, n(%) | 1 (12.5%) | 3 (37.5%) | 0.25014 |
| Relapse, n(%) | 2 (25%) | 1 (12.5%) | 0.52218 |

**Figure 2:**
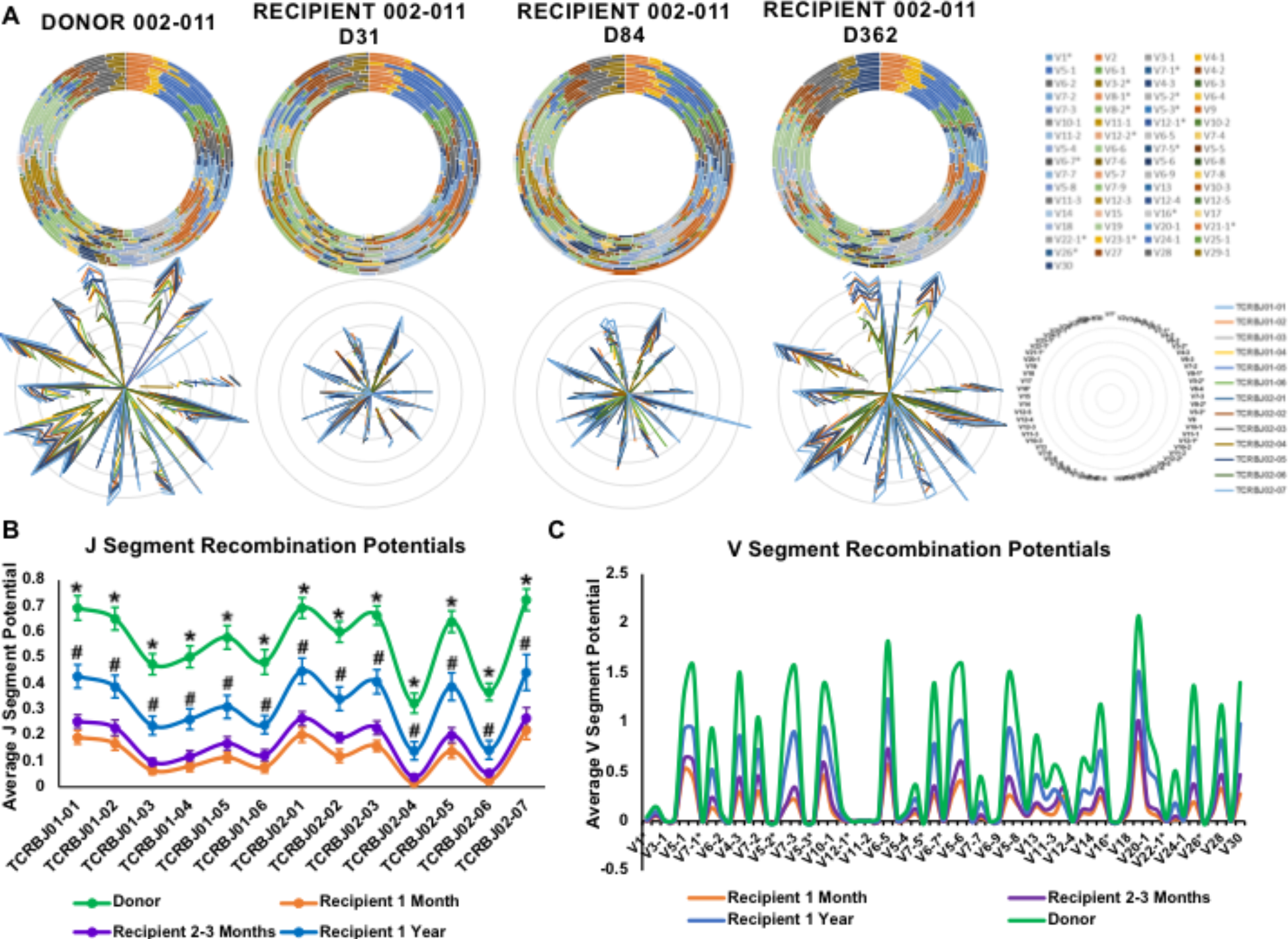
Evolution of the Immune Repertoire after BMT. A) Relative proportional distribution (RPD) graphs (top panel) and radar plots (bottom panel) of a representative bone marrow transplant donor and recipient pair at the indicated days post-transplant. Legends for respective figures are included at far right. B) J segment recombination potential values as derived from the recombination potential equation for donor and recipients at 1 month, 2-3 months, and 1 year post-transplant. * denotes p<0.001, comparing donor versus all recipient time points, # denotes p<0.05, comparing recipient 1 year to recipient 1 month and 2-3 months (n=6). C) Average V segment recombination potentials for donor (n=13) and recipients at 1 month (n=7), 2-3 months (n=7), and 1 year (n=15).

To further explore the self-similar organization of the repertoire and to evaluate for any perturbations after BMT, radar plots were utilized to study the clonal frequency of individual VJ recombinations and assess for the presence of translational symmetry in this process. Natural log-transformed clonal frequencies of TCRs bearing unique VJ recombinations were plotted in the radar plots allowing for direct comparison across multiple recombinations. Translational symmetry was evident across the repertoire when the recombination of each V segment with each of the J segments is examined in the donor (Figure 2A, bottom panel, column 1). In other words, the frequency of each V segment’s recombination with all the J segments is similar relative to its neighboring segments and vice versa. Thus, the recombination frequency of each of the TCR gene segments remains the same across the locus, which manifests as the rings in the radar plots having the same configuration when, for example, all the J segments are analyzed.

Further examination of these recombination frequencies reveals that there is a periodicity to how V segments recombine with each J segment across the locus and vice versa. This periodic frequency distribution and the translational symmetry of VJ recombination is still evident following BMT, and results in preservation of the “frame” of the TCR repertoire after transplant (Figure 2A, bottom panel, columns 2-4). However, the overall complexity of the repertoire diminishes post-transplant, as there is a reduction in the magnitude of the clonal frequency of each of the VJ recombinations plotted, which recovers with time. This is comparable to the difference between a tree in the summer, when it possesses its full accompaniment of leaves, versus in the winter when it is barren. Regardless of season, the basic structure (i.e., the trunk and branches) remains intact, akin to the TCR repertoire where the framework of TRB VJ recombinations is present both pre-and post-transplant, perhaps related to the abundance of these recombinations in the normal setting. This was observed in all donor-recipient samples analyzed (n=9).

The inherent order of VJ recombination observed in the TCR repertoire suggests that the TCR β gene segment usage is not random, either in health or after transplant. Instead, it has been hypothesized that the position of the gene segment on the TRB locus determines the probability of each TCR β V segment to recombine with each of the J segments, to form unique TCR bearing clones.^9^ To investigate this, the average recombination potential for individual J (Figure 2B) or V (Figure 2C) segments was calculated. These calculations encompass both the clonal frequency of VJ recombinations, as well as the distance of the segments under study from the *D* segments of the TRB locus. When these recombination potentials were calculated at various times post-transplant, both J (Figure 2B) and V (Figure 2C) segment recombination potentials demonstrated a periodic, oscillatory patterning of TCR usage from one end of the locus to the other. The symmetry and periodicity with which TCR-VJ recombinations occur here is highly suggestive of a pre-determined rearrangement scheme dictated by gene segment ordering at the genomic level. As seen in the radar plots, translational symmetry in terms of the frequency of VJ recombinations is maintained after transplant (similar pattern of gene segment usage) with complexity (clonal frequency reflected by the magnitude of recombination potential) increasing over time (Figures 2B and 2C). Though recipient J and V segment recombination potentials steadily increase (Figures 2B and 2C; Supplementary Figure 2A), the complexity of the repertoire still doesn’t recover to that of the donor even three years out post-transplant (Supplementary Figures 2B and 2C). This is also reflected in the total unique productive sequences (Supplementary Figure 2E) for the recipients, despite no significant differences in sequencing reads from these samples as compared to donor (Supplementary Figure 2D).

### TCR Repertoire Architecture is Preserved Within Individual T-cell Subsets

Next the order observed in the total CD3^+^ population was evaluated within T cell subsets. CD3^+^CD4^+^ and CD3^+^CD8^+^ sorted cell populations subjected to TCR β locus sequencing demonstrated conserved organizational principles with evidence of selfsimilarity when looking at both RPD graphs (Figure 3A) and symmetry when looking at the radar plots (Figure 3B). This was evident both in donor and recipient CD4^+^ and CD8^+^ T-cell pools at various times post-transplant (Figure 3A and 3B, Supplementary Figure 3A). This supports the hypothesis that the property of self-similar repertoire organization if true for T cells as a whole, may extend to all T cell subsets.

**Figure 3:**
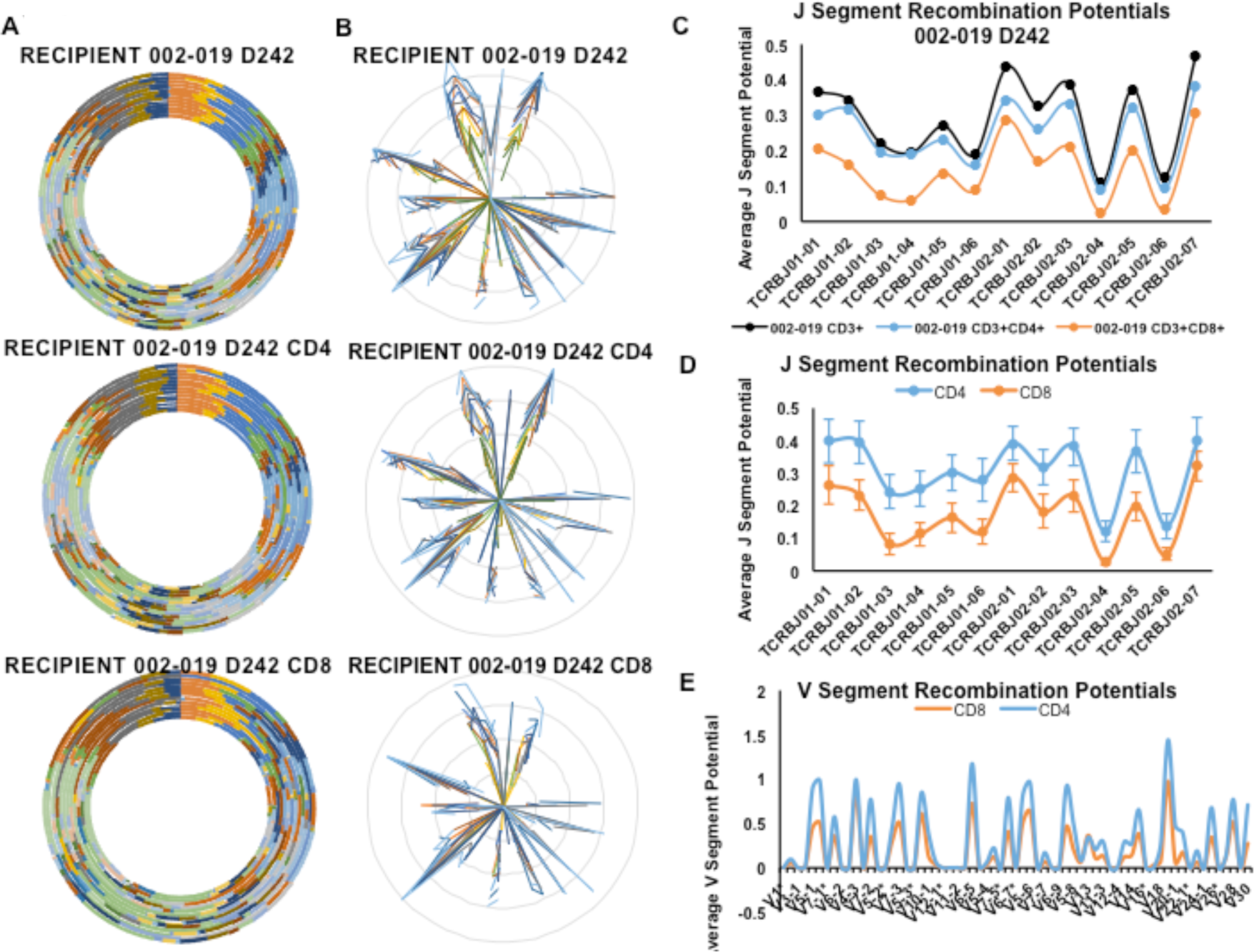
TCR Repertoire Architecture is Preserved within T-cell subsets. RPD graphs (A) and radar plots (B) of unfractionated (top), CD4+, and CD8+ T cell pools in a BMT recipient at day (D) 242 post-transplant. (C) Corresponding J segment recombination potentials for the unsorted CD3^+^ population as compared to CD3^+^CD4^+^ or CD3^+^CD8^+^ subsets. Average J (D) or V (E) segment recombination potentials for CD4^+^ (n=5) or CD8^+^ (n=5) T cells.

The preservation of translational symmetry in terms of TCR usage was also appreciable when evaluating the recombination potential for a representative patient (002-019) at day +242 after transplant (Figure 3C) as the waveform was unaltered for both CD3^+^CD4^+^ and CD3^+^CD8^+^ cells as compared to the total CD3+ population. As was suggested in the radar plots (Figure 3B, Supplementary Figure 3A), complexity of the CD8^+^ repertoire was diminished as compared to that of the CD4^+^ pool (Figure 3C). This was reflected in both the J (Figure 3D) and V (Figure 3E) segment recombination potential values with CD8^+^ T cells consistently lagging behind the CD4^+^ T cells regardless of which gene segment was being interrogated. This methodology recapitulates prior reports demonstrating greater diversity in the CD4^+^ T-cell population.^6,28^ It also suggests that the reduction seen at the CD8^+^ level is not at the expense of TCR VJ recombination events and overall T cell repertoire composition is maintained. This may also be related to differences in CD4 and CD8 structure that affects TCR avidity and results in differences in shaping of the repertoire.^29^

### Organization of Repertoire is Maintained in Patients that Develop GVHD

Acute graft versus host disease (aGVHD) has previously been associated with changes in the TCR repertoire when analyzed via high throughput sequencing. ^7,13,14,16^ We investigated whether or not there was any effect of aGVHD on the organization of the TCR repertoire and its evolution over time. Patients that developed aGVHD did not display any obvious distortion of the previously observed symmetry or the time-dependent increase in repertoire complexity when TCR β sequencing data was evaluated with RPD graphs or radar plots (Figure 4A and 4B). Though there is a suggestion of aGVHD, as well as conceivably its accompanying immunosuppressive treatment, limiting the complexity of the developing repertoire, there was no statistically significant difference observed between recombination potentials between patients when stratified by aGVHD status. Acute GVHD did not affect the translational symmetry of the repertoire observed in the recombination potential plots at either 2-3 months (Figure 4C, middle panel) or at 1 year (Figure 4C, lower panel) post-transplant.This indicates that the representation of various VDJ recombinations in the T cell repertoire is preserved following transplantation, and that either aGVHD is mediated by a polyclonal expansion of pathogenic T cells or that aGVHD is not associated with numerical expansions of pathogenic clones in the blood.

**Figure 4:**
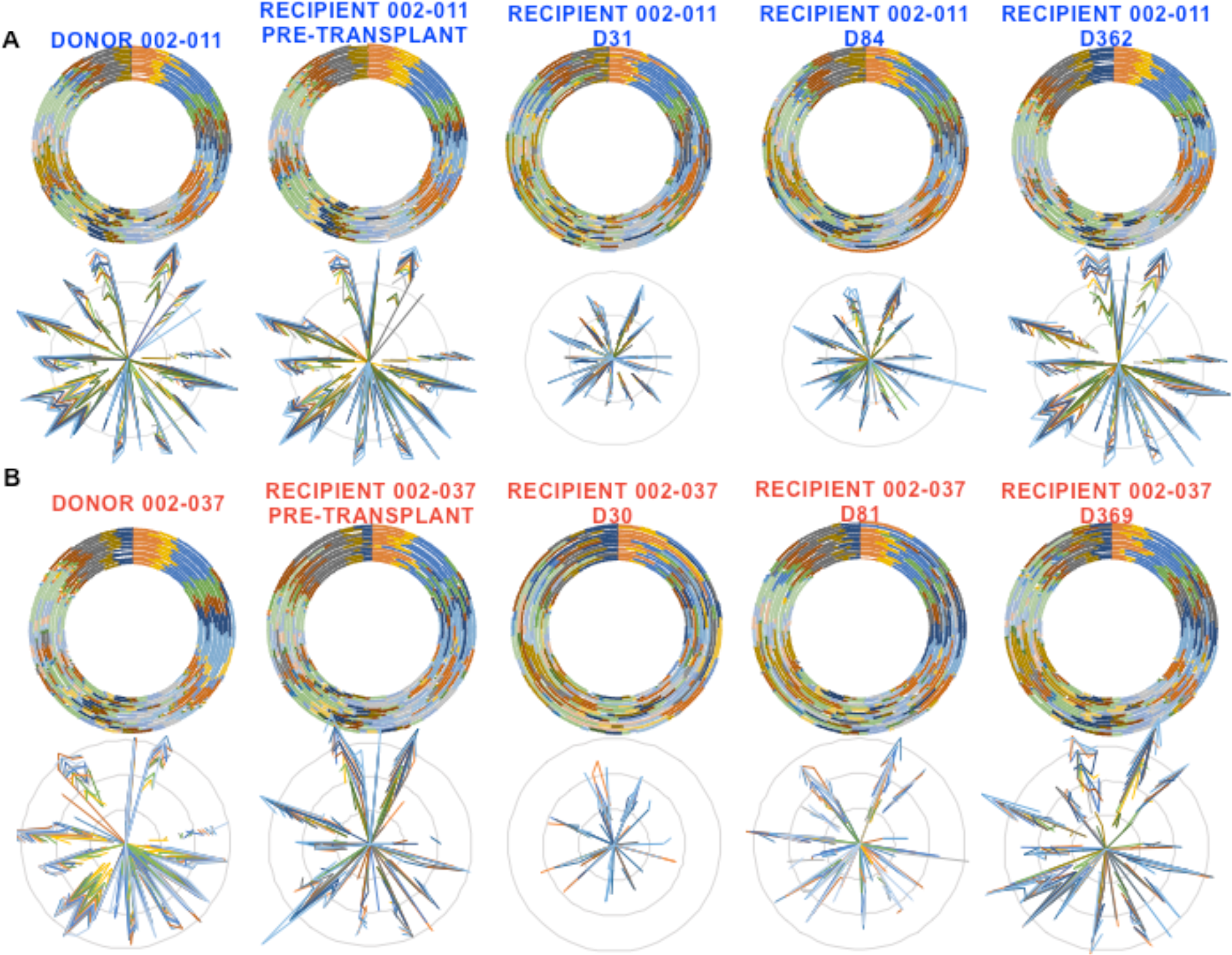
Repertoire Organization is Maintained in Patients who Develop aGVHD. BMT donor-recipient pairs at indicated time with RPD graphs (top) and radar plots (bottom) in a representative patients (A) without GVHD (blue) or (B) with aGVHD (red) demonstrating comparable reconstitution of the immune repertoire. (C) J segment recombination potential values for donors (top) or BMT recipients at 2-3 months (middle) or 1 year post-transplant (bottom) stratified by aGVHD status. Recipients not developing aGVHD are represented in blue, with those developing aGVHD in red. (D) Box and whisker representation of recipient 1-year J segment recombination potential values for patients without or with aGVHD across all J segments (n=7 for GVHD and n=8 for GVHD plots). (E) Average 1-year V segment recombination potential values for patients with (n=7) or without (n=8) aGVHD with donor (n=13) curve included as control.

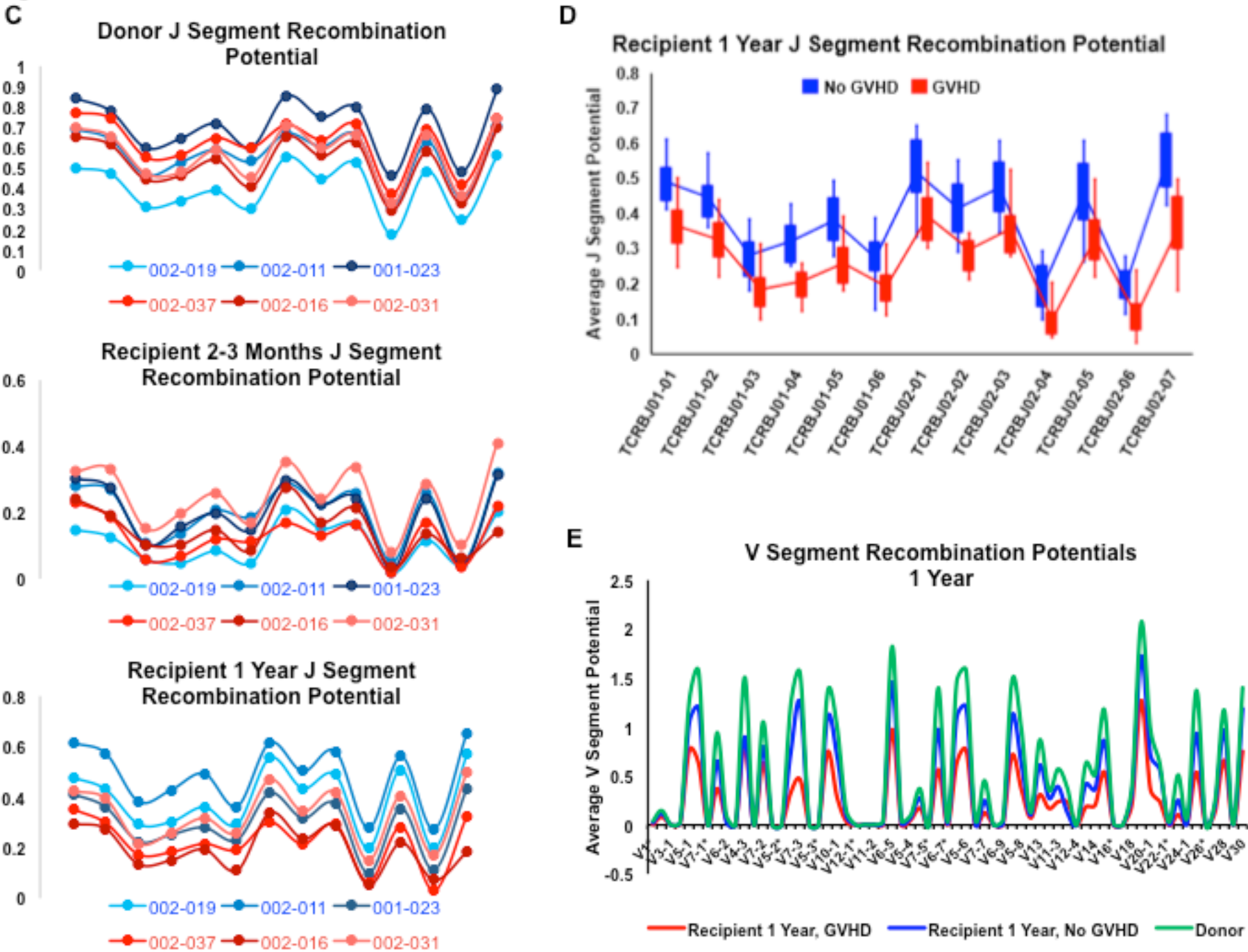

TRB-J (Figure 4C, bottom panel and Figure 4D) and V (Figure 4E) segment recombination potentials at one-year post-transplant demonstrate a trend of lower complexity (i.e., clonal frequency/magnitude) in patients who had developed aGVHD, which we expect is a consequence of both the immunosuppressive regimens administered to treat the GVHD as well as GVHD itself altering T cell clonal expansion patterns. This difference is less evident when repertoires of aGVHD patients are compared to those not affected by aGVHD via RPD graphs or radar plots (Supplementary Figures 4A and B). Further, there was no appreciable difference in repertoire complexity when J or V segment recombination potentials are calculated at early times posttransplant in cases of aGVHD (Figure 4C, middle panel and Supplementary Figure 4E and F).

When evaluating differences in the donor repertoires, there are no obvious differences in overall repertoire organization or complexity that may be predictive of recipient tendency to develop aGVHD (Figure 4C and Supplementary Figure 4G). There was also no difference in the total CD3 counts or absolute lymphocyte counts in those patients that developed aGVHD (Supplementary Figures 5A and 5B) that might account for the differences observed in recombination potential values at one year. There was a trend towards reduced total sequencing reads and unique productive sequences in GVHD patients only at 1 year possibly suggestive of diminished overall T cell reconstitution likely exacerbated by immunosuppressive therapies in those patients. The diversity indices between patients with and without aGVHD however, showed no significant differences (Supplementary Figures 5E and 5F).

Prior studies have noted that CMV serostatus and reactivation are associated with an earlier recovery in lymphocyte counts with a shift towards an increasingly clonal and less diverse repertoire.^8,30^ In examining the organization of the repertoire in BMT recipients with either positive or negative CMV serostatus, both J and V segment recombination potentials showed similar results (Supplementary Figures 4C and 4D). Together, these findings point to maintenance of repertoire architecture and its underlying determining principles in both health and disease states alike. It is important to recognize the impact that these calculations have on scaling of T cell clonal frequencies. Log transformation of the T cell clonal frequencies may obscure relative clonal expansion, which may occur in the event of a specific infection such as CMV reactivation or GVHD. The maintenance of overall clonal architecture in this instance is important in that it provides the basis for eventual recovery of the T cell repertoire, given adequate stimulation with appropriate antigens, for example as in responses observed to vaccination or actual infections.

### Low Frequency Donor Clones May Mediate GVHD

With the emergence of high throughput sequencing techniques, there has been a great deal of interest in trying to determine if particular TCR clones can be linked to GVHD, or alternatively if there are significant shifts in the overall repertoire make-up due to oligoclonality. Previous work had demonstrated that the self-similar distribution and organization of the repertoire could be evaluated through log transformed data to determine the fractal order ^22,31^ of the T cell population being studied. Consistent with the results from the analysis of the RPD graphs, radar plots, and symmetry calculations, there was no significant effect on the clonal hierarchy in patients that developed aGVHD as assessed by log-rank plots (Figure 5A). Slopes of these plots showed near uniformity reflective of the preserved self-similar nature, with Power Law clonal frequency distribution of the T cell pool in patients both with and without GVHD, regardless of post-transplant time (Table 2). Further, when the percentage with which each rank constituting the overall T cell population was assessed, patients with aGVHD did not appear to have any distortion in their overall representation in the top ranks, with only observable differences in the way the lower ranked clones were distributed (Figure 5B, note ranks 5 and 6 at 1 month and 2-3 months post-transplant). Similar to the symmetry analysis, these findings suggest that GVHD may be a polyclonal process with large numbers of T cell clones being involved in its pathogenesis, as opposed to an oligoclonal proliferation with a limited number of dominant T cell clones. Alternatively, significant oligoclonal expansion may predominate in the target tissues of GVHD that may not be detectable at the same level in peripheral blood.

**Figure 5:**
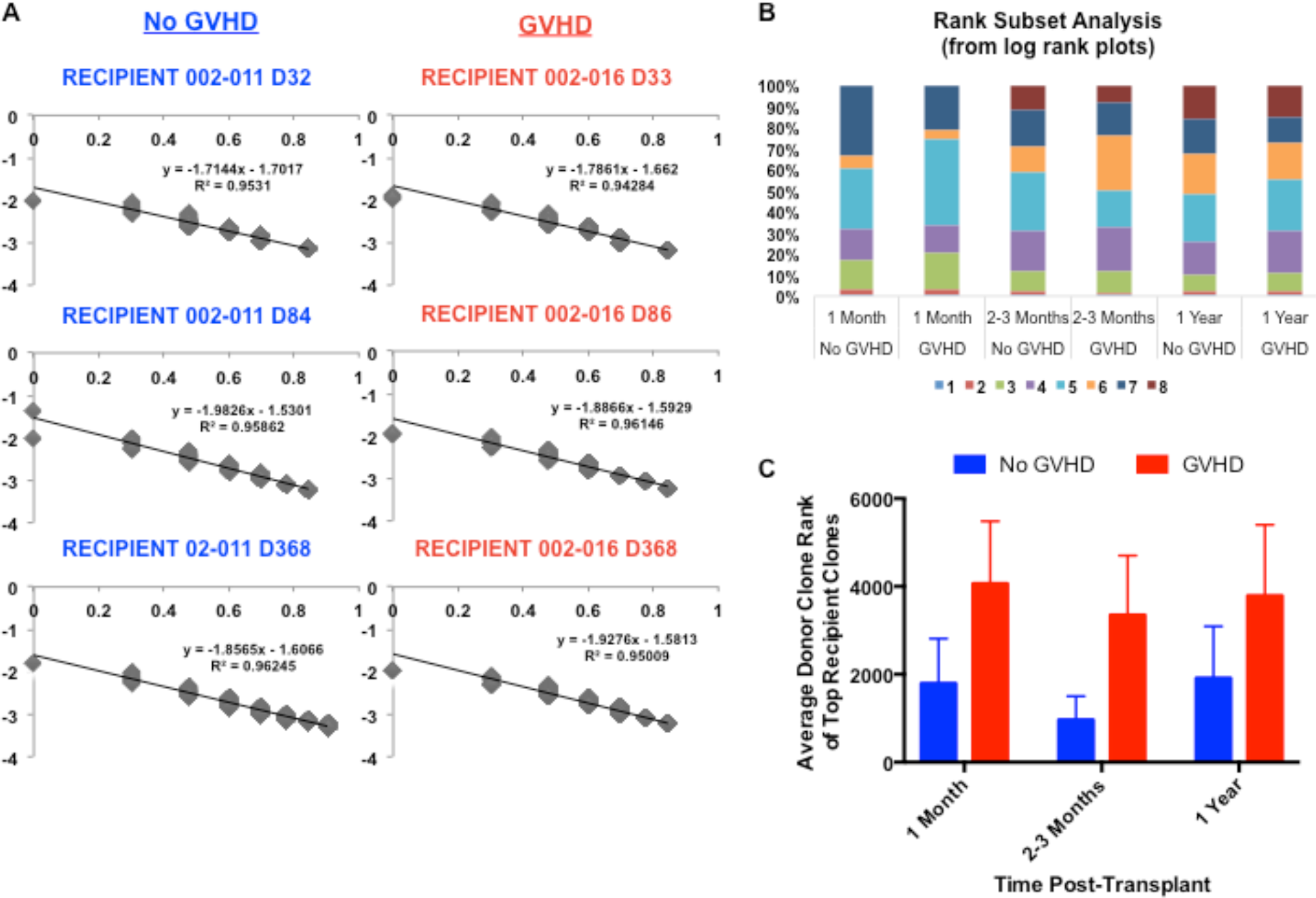
Self-similarity and Clonal Hierarchy are Preserved in aGVHD. (A) Log-log rank plots of representative BMT recipients without (left, blue) or with (right, red) aGVHD at the indicated day post-transplant. Linear regression analysis is included in each plot where the X axis equals log (relative clonal frequency) and the Y axis represents log(clone rank). (B) Rank subset analysis for patients without (n=4) or with (n=3) aGVHD depicting overall percent contribution of each rank to the repertoire at the indicated times post-transplant. TCR clones were ranked based on their clonal frequency according to the rules for the log-log rank plots from 1 (highest frequency) through 8 (lowest frequency). (C) Average donor rank of the top 5 recipient recombination events (based on productive frequency and shared with donor) in patients with (n=4) or without (n=4) aGVHD.

**Table 2:** Self-similarity and fractal order are not distorted in aGVHD Patients. Absolute value of log-log rank plot slopes in patients with (n=3) or without (n=3) aGVHD at 1 month, 2-3 months, or 1 year post-transplant.

| No GVHD |  |  |  | GVHD |  |  |  |
| --- | --- | --- | --- | --- | --- | --- | --- |
|  | 1 Month | 2-3 Months | 1 Year |  | 1 Month | 2-3 Months | 1 Year |
| **002-019** | 1.760 | 1.774 | 1.807 | **002-037** | 1.576 | 1.789 | 1.805 |
| **002-011** | 1.714 | 1.983 | 1.857 | **002-016** | 1.786 | 1.887 | 1.928 |
| **001-023** | 1.675 | 1.836 | 1.734 | **002-031** | 1.743 | 1.720 | 1.742 |
| **Average** | 1.717 | 1.864 | 1.799 |  | 1.702 | 1.799 | 1.825 |
| **(±SEM)** | (0.025) | (0.062) | (0.035) |  | (0.064) | (0.048) | (0.054) |

Prior reports have implicated the possibility that there might be pathogenic, infused donor T cell clones that mediate aGVHD.^8^ Given the relative limited functionality of the thymus post-transplant as reflected by low numbers of recent thymic emigrants, ^8^ it is likely that T cells mediating aGVHD may be present in the donor product and proliferate upon antigen encounter in the recipient. These maladaptive T cell clones would likely be present at a low frequency in the donor, prior to proliferating and becoming dominant mediators of alloreactivity in the recipient. When evaluating those T cells clones that constituted the top recipient ranks (i.e. highest frequency clones post-transplant), patients that developed aGVHD tended to have a larger proportion of T cell clones that were present in the donor at a low frequency as compared to the patients who did not develop aGVHD (Figure 5C). Reciprocally, significantly fewer of the highest frequency donor infused T cell clones were found in recipients that developed aGVHD (Figure 6A), particularly in the early post-transplant period, coinciding with the time frame of aGVHD pathogenesis. This supports the notion that lower ranked donor T cell clones proliferate to a greater extent in patients with aGVHD, likely upon encountering alloreactive antigens not present in the donor. This is consistent with the notion of T cell ‘vector transformation’ post-transplant, recently proposed as part of the dynamical system theory of T cell response. ^20,32^ In agreement with the observations from the RPD graphs, radar plots, and recombination potential calculations, there was no increase in repertoire clonality (based on (1-entropy) and Gini coefficient calculations) in patients that developed aGVHD (Figure 6B and 6C). There was also no difference in donor repertoires when stratified by recipient GVHD status (Supplementary Figures 6A and 6B). This implies that while a full complement of T cell clones may be observed in patients with aGVHD following posttransplant cyclophosphamide based GVHD prophylaxis, there is a significant shift of the relative rank of T cell clones from the donor. Taken together these results imply that aGVHD may be driven by multiple alloreactive, pathogenic T cell clones in the infused donor product.

**Figure 6:**
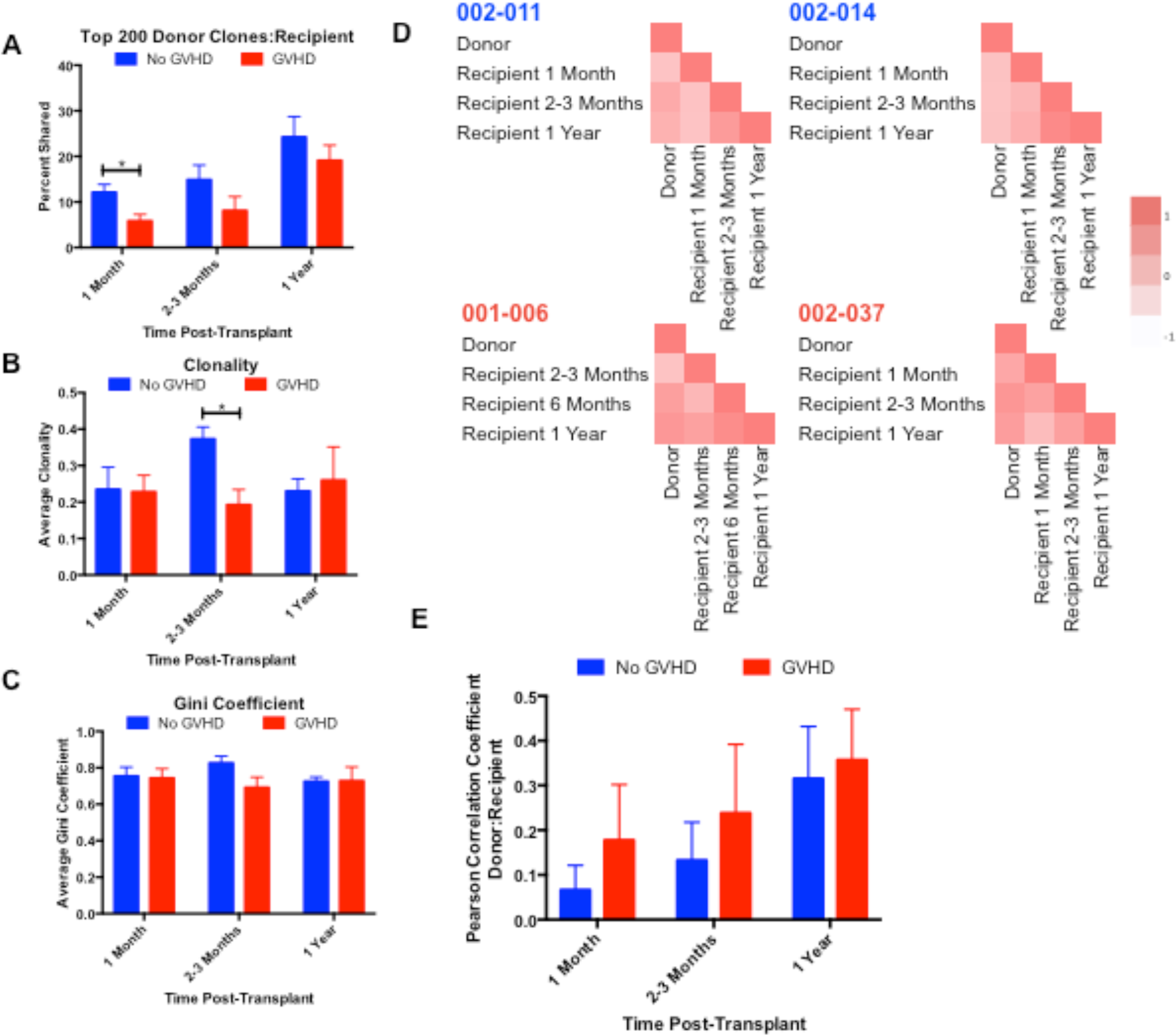
Implications of Donor-Recipient T cell Clonotype Disparities and Dynamic changes Post-transplant. (A) Percent of top 200 donor clones expressed in the corresponding recipient rank at the indicated time points post-transplant in patients with (n=4) or without (n=4) GVHD. * denotes p<0.04. Clonality (B) as defined by 1-entropy (from Shannon Weaver entropy equation) and Gini coefficient (C) values in BMT recipients at 1 month (n=4), 2-3 months (n=4-5), or 1 year (n=4) after BMT separated based on presence or absence of aGVHD. * denotes p=0.01. (D) Representative heat maps of Pearson correlation coefficients comparing TCR β rearrangement relatedness of donor:recipient and intra-recipient time points post-transplant in patients without (top, blue) or with (bottom, red) aGVHD. (E) Quantitative analysis of plots in (D) depicting the average Pearson correlation coefficient comparing donor T cell recombination events to recipients at indicated times post-transplant in patients with (n=4) versus without (n=4) aGVHD. (F) Representative heat maps of VJ recombination matrices demonstrating percent change between donor and recipient at 1 month, 2-3 months, or 1 year posttransplant across all potential VJ recombination events. Recipient with GVHD is depicted in the right panels. (n=5), Green=decreased, Red=increased (G) Intra-recipient VJ recombination matrices demonstrating percent change in VJ recombination events within the indicated recipients over-time (n=5).

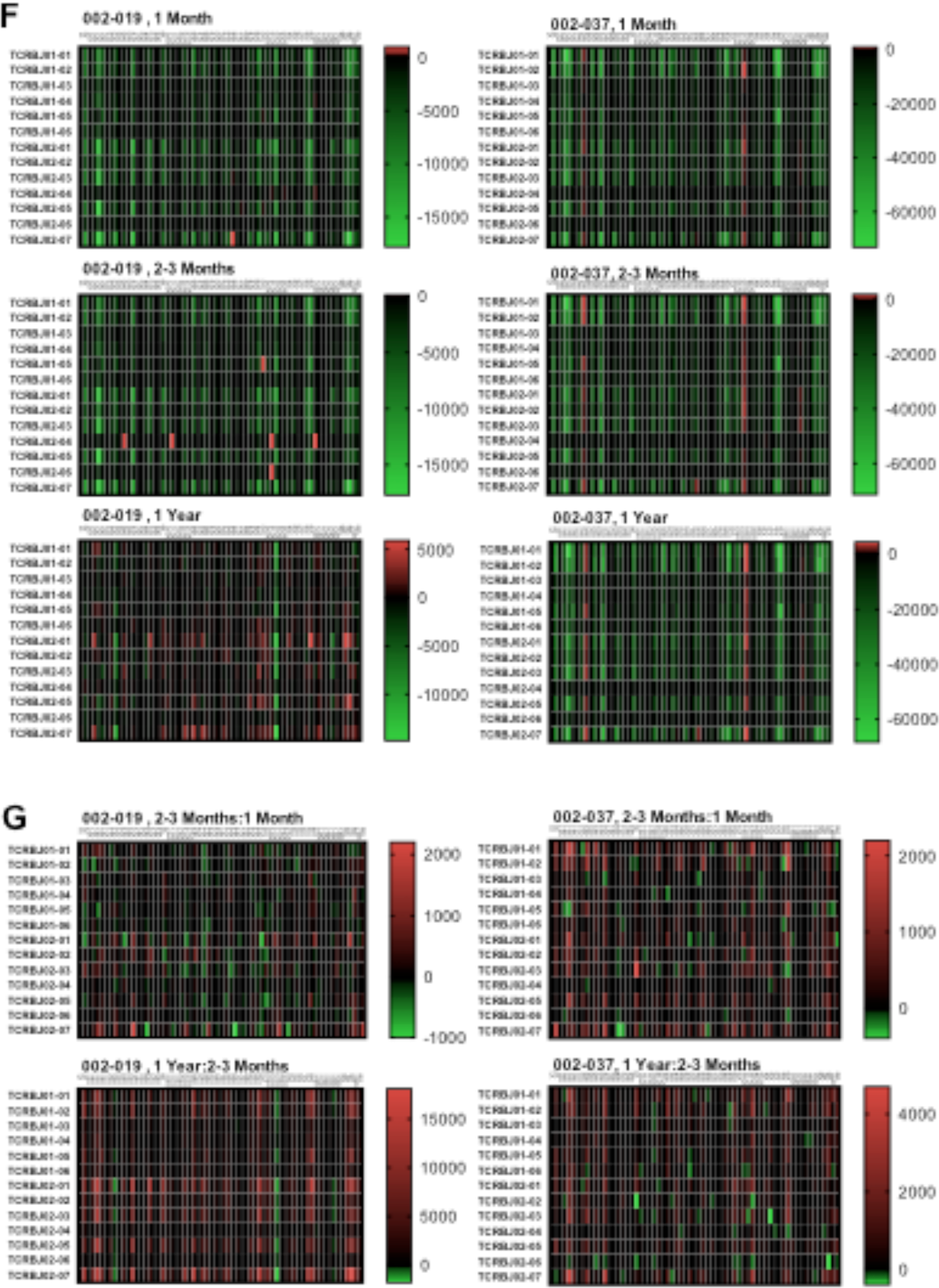

When CMV serostatus was evaluated as to its effect on the developing T cell repertoire, consistent with prior published reports^8^, a greater overlap was observed between donor and recipient repertoires in recipients who had a positive CMV serostatus using this methodology (Supplementary Figures 7A, 7B, and 7E). This was accompanied by increased clonality in the repertoire over time likely reflective of T cell clonal amplification by CMV antigens (Supplementary Figures 7C and 7D) as previously described.^8^.

Overall, a dynamic and continuously evolving T cell repertoire is observed following BMT, which makes clonal tracking and monitoring of clonal frequency changes in the post-transplant setting extremely important to understand the cellular basis of alloreactivity. Regardless of GVHD status, there is a dramatic change in the recipient repertoire as compared to the donor repertoire when the Pearson correlation coefficient is calculated for donor-recipient pairs post-transplant (Figure 6D and 6E). This is also evident when comparing donor to recipient (Figure 6F) or intra-recipient (Figure 6G) repertoires over time using V × J matrices to depict T cell clones. These depict the percent change for individual VDJ recombinations between the repertoires at different times. In both cases, there is an obvious shift in the repertoire hierarchy. This highlights, the importance of thoroughly quantifying the donor immune response in the post-transplant setting. As demonstrated here this may be accomplished by mathematically characterizing the post transplant T cell repertoire evolution. This may potentially predict transplant outcomes or help us understand the impact of interventions, such as vaccines, in the post-transplant setting.

## Discussion

The human T cell repertoire is complex and comprised of cells with different functional roles, providing comprehensive protection from a legion of microbes encountered in daily life. This requires recognition of the myriad antigens these microbes harbor, a task accomplished in collaboration with the cells of the antigen presenting and processing system. Antigens presented by dendritic cells, macrophages, and other antigen presenting cells are recognized by T cells by means of T cell receptors which can take on an equally large body of configurations as the antigen array encountered by the individual. This comprehensive, yet precise ability to recognize antigens and react, while beneficial under normal circumstances, poses a significant barrier to both solid organ and bone marrow cell transplantation. The donor T cell response to recipient antigens, presented on matched or mismatched HLA molecules can be devastating and lead to tissue injury involving many different organ systems, termed graft vs. host disease (GVHD). Over the years much effort has been expended on trying to understand the changes brought about in the T cell repertoire of patients who develop GVHD as compared to those who do not, to enable better early identification and possibly preemptive therapy to be instituted.

The T cell receptors are comprised of α and β subunits, which each are made up of variable and joining, as well as diversity segments. Their recombination, and the addition of non-template, nucleotides confers the clonal diversity on the T cell repertoire, enabling it to respond to changes in the antigenic landscape. This is a dynamic process, with T cell clonal hierarchy changing in response to the different antigens encountered. As such, following BMT where there is a completely new library of recipient minor histocompatibility antigens that donor T cells may encounter, the clonal hierarchy is likely to change very significantly from the steady state present at the time of stem cell donation.^20^ This may also be the case for major histocompatibility antigens in the context of haploidentical transplantation. This was corroborated by the finding of a change in the clonal hierarchy reported here, in that T cell clones that were dominant in the donor declined after transplant, and others which were present in low numbers proliferated in the recipient. It is also noteworthy that there was significant dynamic change seen over time in the clonal makeup of the recipient when the T cell repertoire was compared over time following BMT. These findings support the dynamical system theory of immune response which posits that antigen driven T cell proliferation may be computed using logistic equations, accounting for antigen affinity for HLA and TCR.^19^ This would imply that T cell responses occur with mathematical precision and are thus susceptible to computation and simulation. Indeed, a methodology for such simulations has been developed which utilizes antigen arrays and computes the ‘vector’ transformation that responding T cells will undergo.^20,32^

If the T cell antigen responses are subject to mathematical rules then, logically the process of T cell receptor generation itself may be so founded. Processes governed by mathematical rules will yield quantifiable results: in other words, the T cell repertoire organization cannot be random. T cell receptor sequencing allows an evaluation of the T cell repertoire by quantifying the contribution of each gene segment to the receptor makeup of the T cell clones. When viewed from the point of view of gene segment usage in the T cell population, fractal organization with self-similarity has been demonstrated previously.^22,31^ This means that when the T cell clonal frequencies are measured using either J segment usage, or VJ segment usage and beyond, similar but not identical clonal distribution is observed at each organizational level.^22^ This fractal organization has also been observed at the level of T cell receptor locus.^9^ Supporting this concept it has also become increasingly recognized that T cell repertoires amongst individuals are remarkably well conserved,^33,34^ and may be more alike at birth with divergence with age due to antigen stimulation.^35^ A property of self-similar systems is that they demonstrate symmetry. Symmetry is a property that is most easily discerned in physical objects, such as the human body, which demonstrates reflection symmetry (appears the same bilaterally), but less obvious in objects that do not have a geometrically definable form. In the case of the T cell repertoire, if the process of VDJ recombination is viewed with reference to the positions of the V and J segment positions on the TCR locus, to identify the symmetry at work in immune repertoire generation. This spatial symmetry is referred to as translational symmetry (Figure 1), which for the VDJ recombination process implies that each J segment will recombine with a particular V segment in proportion to its position on the locus and vice versa. This is evident in the radar plots depicted in this paper which plot the clonal frequencies of various recombinations which show concentric rings of V segment defined clonal frequency when specific J segment containing clones are plotted. It is to be noted that these plots are on a log scale, which is a consequence of the exponential growth of the T cell clones in response to their cognate antigens. That, translational symmetry of VDJ recombination is present in the normal and post-transplant T cell repertoire, with diminished complexity of the repertoire, supports the application of quantitative reasoning to more accurately measure broad T cell responses following various methods of immune suppression after BMT.

While translational symmetry of TRB VDJ recombination (and logically TRA recombination) is relatively well preserved, the repertoire complexity is not. This is likely a consequence of the kinetics of adult lymphoid repopulation after immunoablative therapy in a post-thymic individual, but more significantly may be an effect of the immunosuppressive therapy following BMT. This effect was manifest in the loss of magnitude in both the symmetry radar plots as well as the T cell receptor recombination-potential calculations. This complexity was restored over time after transplant, likely due to both individual clonal growth as well as increasing clonal diversity. The inference of increasing clonal diversity is supported by the fact that translational symmetry of VDJ recombination is maintained through this process. It is noteworthy that this set of measurements is not dramatically impacted by either GVHD, or CMV serostatus. It should however be noted that, while these analyses account for the scaling bias introduced by unique clonal growth at the time of measurement and make it possible to compare measurements across several points in time, the isolated clonal T cell expansion occurring in response to unique antigens may be obscured.

The consistent TCR usage observed within these studies amongst BMT donors and recipients, regardless of post-transplant time, points to an ingrained order to which VDJ recombination occurs. One potential confounder of the above results noted in the literature is that these may be the byproduct of PCR amplification bias. However, through the use of primer optimized multiplex PCR, the sequencing platform utilized to generate these results can minimize bias through computational corrections.^26^ Moreover, the observations reported here are remarkably similar to those of a prior model of J segment rearrangement biases, which utilized a mathematical model to faithfully predict J segment usage in a murine model.^23^ In that report they had also utilized transcript data from antigen experienced human T cells, as well as CD4^+^ and CD8^+^ cells, to try and determine the relative frequencies of J segment usage in humans. The high-throughput sequencing results here are similar to those frequencies, indicating that gene segment usage both at the TRB V and J loci follows an inherent pattern of recombination, and is not just an artifact of methodology. What’s more, this is appreciated even early on after BMT and evolves in this manner throughout the posttransplant period in all donors and recipients studied. The driving force behind this is likely the organization of the DNA itself,^9,36,37^ whereby its physical structure and fractal arrangement may increase the probability of certain VDJ rearrangements occurring and, and thus, contributing more substantially to the immune reconstitution post-transplant. It is conceivable then that this observation extends beyond TCR recombination and is important to consider in a variety of biological processes. It also explains why the overarching architecture seen within the total CD3 population also persists at the CD4^+^ and CD8^+^ subset levels.

It is possible that the relative preservation of symmetry after transplant may reflect relative clonal abundance in the graft or the unique immune suppressive mechanism of post-transplant cyclophosphamide. Further work will be needed to clarify the *de novo* contribution of the thymus in the post-transplant setting, which is generally thought to be limited by age of transplant recipients, conditioning regimen, and disease state, particularly in the context of GVHD.^38,39^ This is especially relevant given the relative low numbers of recent thymic emigrants observed after BMT in these largely older patients (median age, 54.5) treated with myeloablative conditioning,^8^ and the role of the thymus in establishing TCR diversity, even in the face of a potentially restricted donor repertoire. ^40^ However, the persistence of the TCR repertoire order over time, and in several different individuals, makes it plausible that this reflects a true underlying quantitative principle of the TCR rearrangement process. Does this have any clinical significance or evolutionary advantage? The obvious answer will be that it provides a robust foundation of receptor diversity on which the T cell shield for pathogens may be built. This may be likened to the analogy of the difference in the shade provided by a tree with its foliage in summer, as opposed to that provided by one bereft of its leaves in the winter. This finding also suggests that patients with preserved repertoire symmetry after they have recovered from GVHD or other insults, can then go on to recover normal immune function and may benefit from interventions such as, vaccinations over time to more fully reconstitute their T cell repertoire.

The methodology presented herein of analyzing the TCR repertoire provides a novel means of comprehensively analyzing the entire repertoire and the dynamics of its evolution over time while accounting for its inherent complexity. The relative preservation of the repertoire architecture in spite of maladaptive processes like aGVHD is counterintuitive, but not completely unexpected, given that GVHD has been shown to affect repertoire complexity across the full spectrum^6,7,14,16^ when classical measures such as clonality and repertoire diversity are utilized for comparisons sake. Further, the preservation of symmetry implies that pathogenic T cell clones do not have a preference for one TCR VDJ configuration or another. This plurality of potentially pathogenic T cell clones mirrors the very large array of potentially alloreactive peptide antigens bound to HLA molecules in transplant donor-recipient pairs. ^32,41^ Further, the range of T cell clonal frequencies observed here, is similar to the variability in HLA binding affinities of the alloreactive peptides in these pairs. This also suggests that the while T cell growth posttransplant may follow mathematical precision, the likelihood of alloreactivity is still a stochastic function of alloreactive T cell clones present in the infused allograft and the antigens presented. Further, GVHD may be a polyclonal T cell response to antigenic disparity between donors and recipient, and not necessarily an oligoclonal process with limited number of dominant T cell clones. This implies that even a ‘weak’ antigen may trigger a pathological graft vs. host response in a particular tissue. This tissue injury response may then be amplified by other T cell clones which target different alloreactive antigens presented following the initial insult.

Is it possible that these quantitative relationships influenced by the GVHD prophylaxis regimen used? As the patients in this study all were treated with posttransplant cyclophosphamide, there is a possibility that the lack of a significant difference in the T cell repertoire between patients with and without aGVHD could be a consequence of this prophylactic regimen. Despite these limitations, there was evidence of the emergence of lower ranked donor clones becoming dominant in the recipient to a greater extent in patients who developed aGVHD in the present study. This was the case when the top 5 T cell clones were analyzed and corresponding to this the top 200 donor clones contributed <5% of the top recipient clones in patients with aGVHD as opposed to those without. While this effect may diminish if a greater proportion of the repertoire is analyzed in a larger number of patients, it does demonstrate a difference in the T cell clonal hierarchy in patients with aGVHD. The findings reported here also reflect changes in the circulation, and it should be noted that these might not accurately reflect the clonal distribution in the lymph nodes and target tissues. Therefore, correlative analysis with tissue samples ^16,42^ in the case of GVHD, considered along with GVHD predictive models utilizing exome sequencing data to elucidate the antigenic background in the context of specific HLA may help to develop an understanding of GVHD pathophysiology at a cellular level and to better predict those patients that might at risk for GVHD. ^20,32,41,43^ With increasing knowledge about the complexity of the immune environment and the role that other T cell subsets play in the pathogenesis of aGVHD, particularly regulatory T cells^44^, it will be important to see whether or not there is any distortion in the TCR architecture of those subsets that may be associated with clinical outcomes such as GVHD.

Above limitations aside, this analysis describes the presence of several tangible physical qualities of the T cell repertoire when viewed from the TRB VJ recombination perspective. These are fractal organization with self-similarity, symmetry, and another mathematically definable quality, the periodicity of the T cell clonal frequency across the TRB locus. This has been shown previously,^9^ and it is to be noted that this periodicity occurs at a scale, which is vastly greater than the size of histone subunits comprising the TRB locus. The periodic nature of VDJ recombination is borne out by other analyses of T cell recombination. It is also noteworthy that the resulting clonal frequencies are scaled logarithmically, as is the TRB locus. These fundamental properties of the T cell repertoire are strong evidence that the observations reported here are not the result of random experimental error, rather they reflect the nature of the TCR gene locus and the rules governing its recombination. Considering the fractal organization with translational symmetry observed in the recombination process, along with the log-periodic nature of the T cell receptor locus, an inescapable conclusion is that nucleotides on TRB locus, and by extension on DNA molecules in general, usually thought of as numbers on a real number line (+ or − numbers), may also be considered as positions in a complex number plane. Complex numbers have a real component as well as an imaginary component, taking the form *x* + *a*√−1 and are traditionally depicted as coordinates in a plane, where the real number component spans the x-axis and the imaginary number component spans the y-axis. Complex numbers are linked to real numbers by the famous Euler’s identity, *e*^*iπ*^ + 1 = 0, where *i* = √−1, and are widely used to represent periodic phenomenon.^45^

In conclusion, we report a novel method of analyzing T cell receptor high throughput sequencing data, which accounts for the TRB locus organization and the quantitative principle governing VDJ recombination. This demonstrates that T cell repertoire evolution post-transplant is a process consistent with dynamical system model of immune reconstitution that can be mathematically ascribed, thus opening the door for better predictive models in the post-transplant setting.

## Acknowledgements

AAT was supported, by research funding from the NIH-NCI Cancer Center Support Grant (P30-CA016059; PI: Gordon Ginder, MD). Author Contributions. JM, Developed the idea, collected data, performed analysis and wrote the paper. MF, HA, collected data and performed analysis and wrote the paper. LL, CK, JR and AT developed the idea and wrote the paper. There are no relevant conflicts of interest to disclose.

---

**Supplementary Figure 1:**
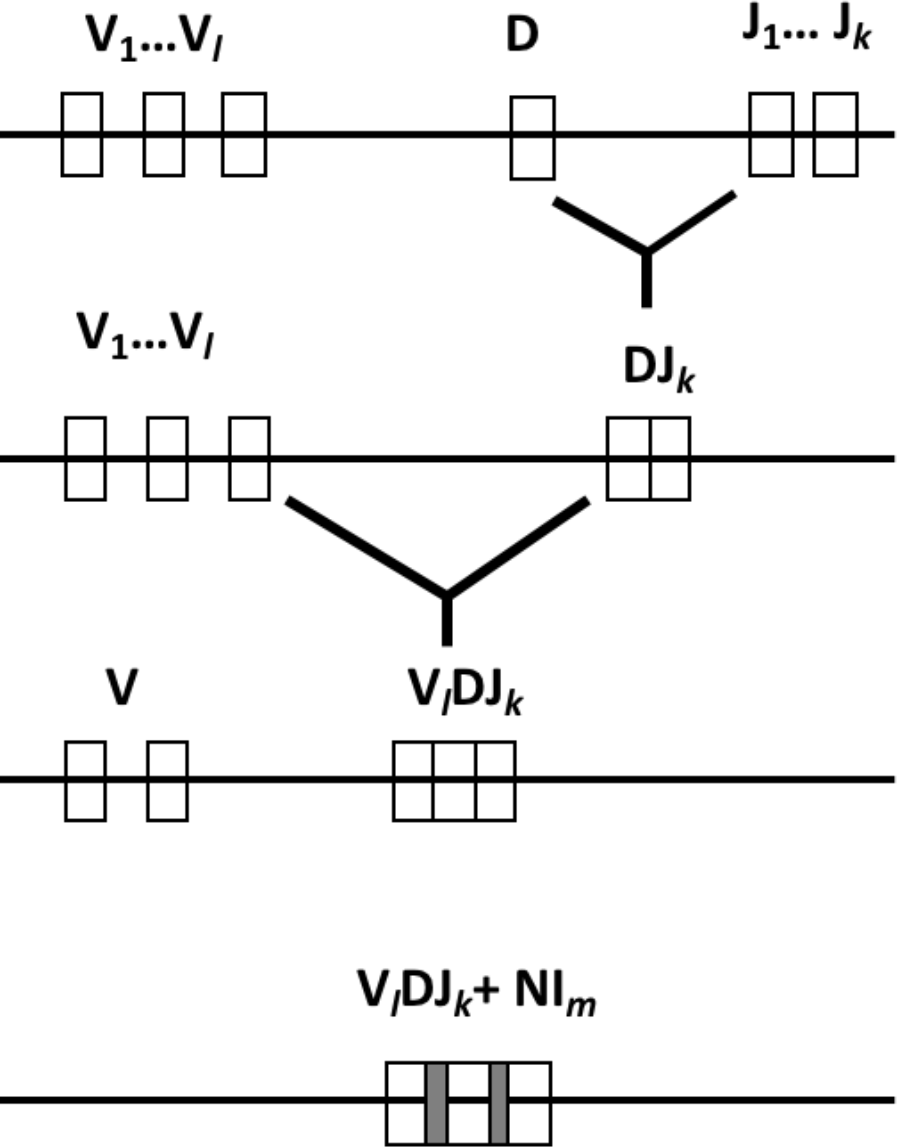
TRB VDJ recombination.

**Supplementary Figure 2:**
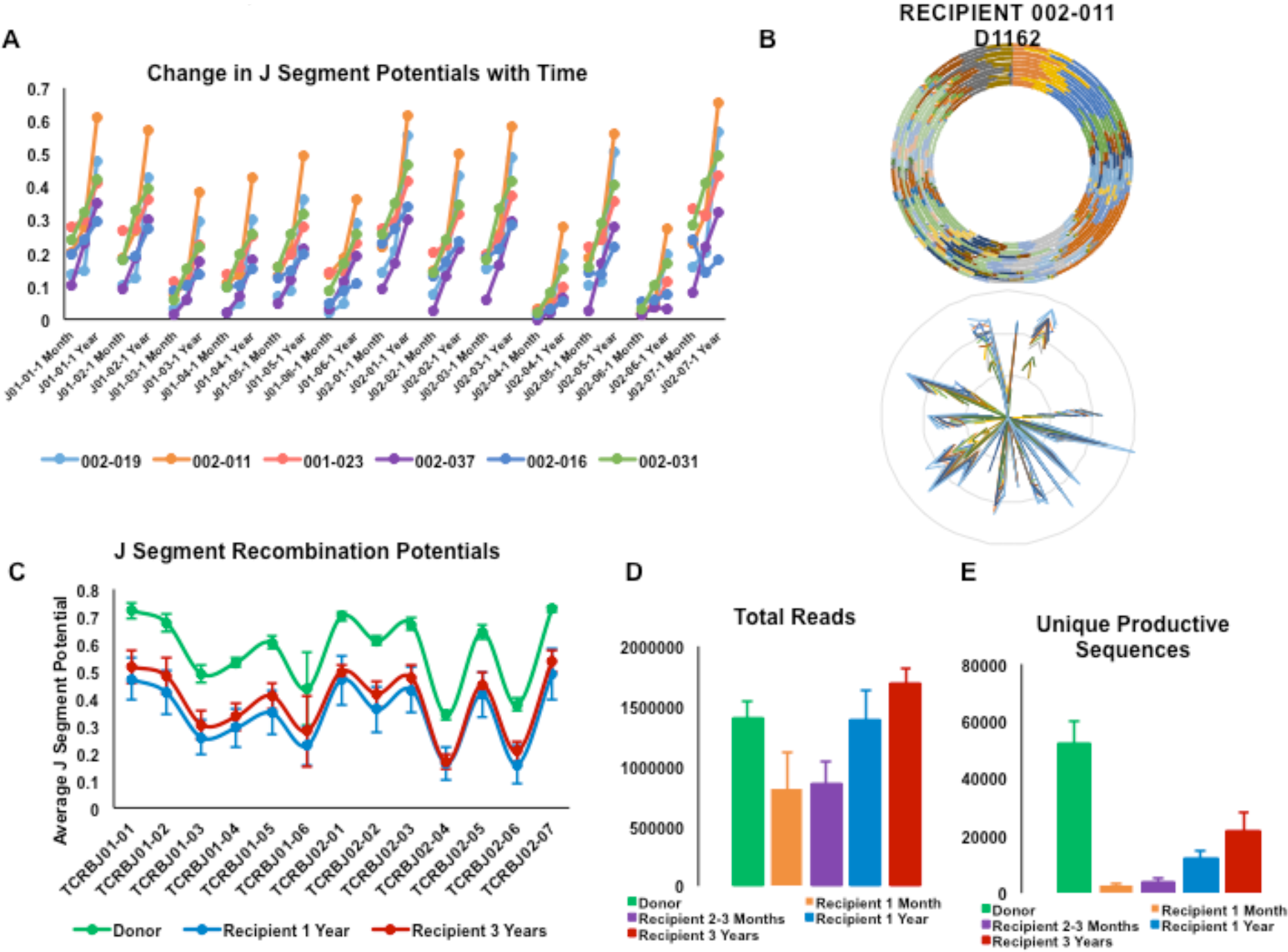
Repertoire Complexity Remains Diminished Years After Transplant. (A) Tracking progression of J segment recombination potential values across each J segment over time for indicated BMT recipients (n=5). (B) Representative RPD graph and radar plot of a recipient 3+ years post-transplant. (C) J segment recombination potentials comparing donor and recipient 1 and 3 years post-transplant (n=3). Average total reads (D) and unique productive sequences (E) from sequencing samples for donor or recipient at the indicated times after BMT.

**Supplementary Figure 3:**
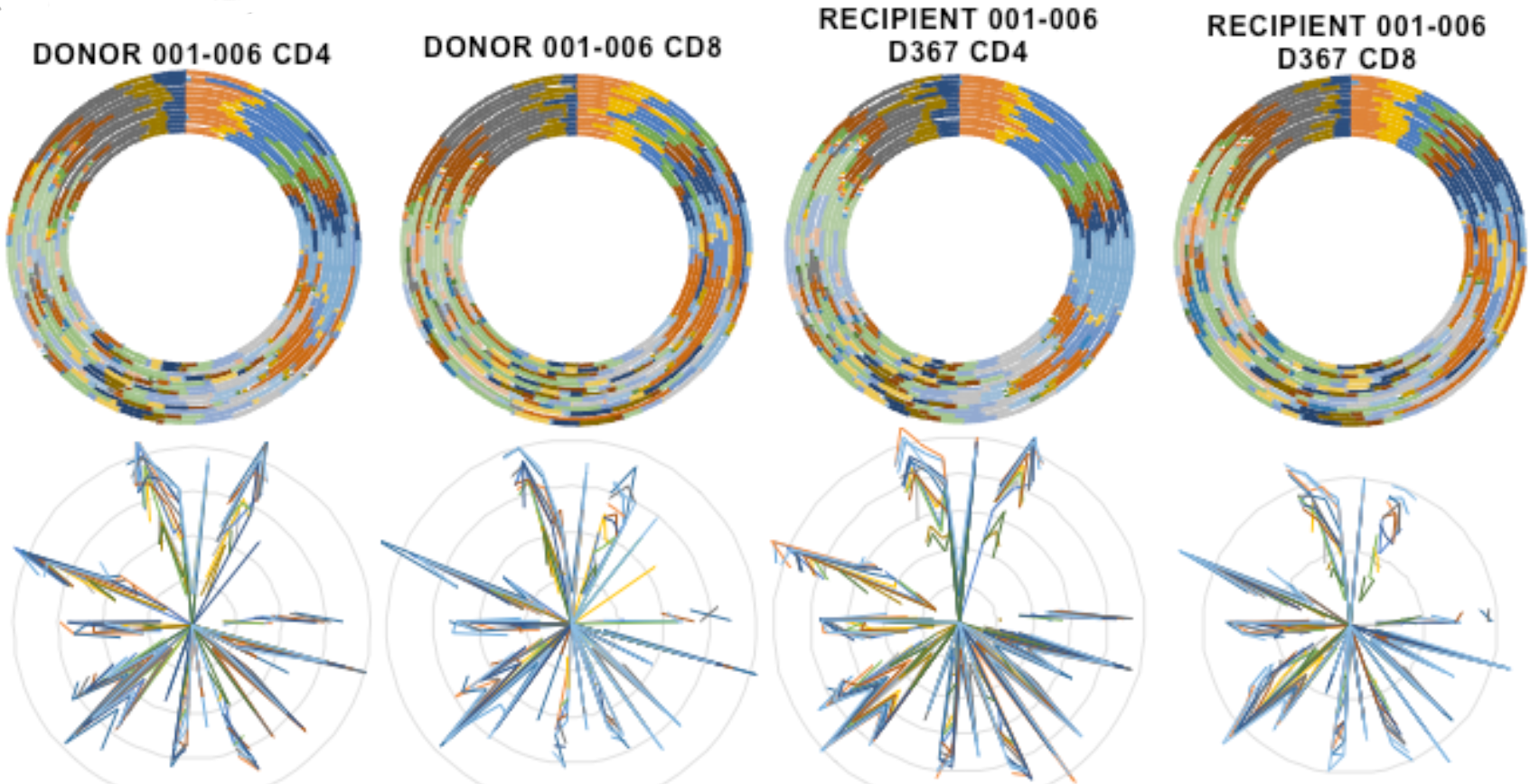
Maintenance of Symmetry in T cell Subsets. RPD graphs (top) and radar plots (bottom) of CD4 and CD8 TCR β VJ recombination events in a BMT donor and recipient pair.

**Supplementary Figure 4:**
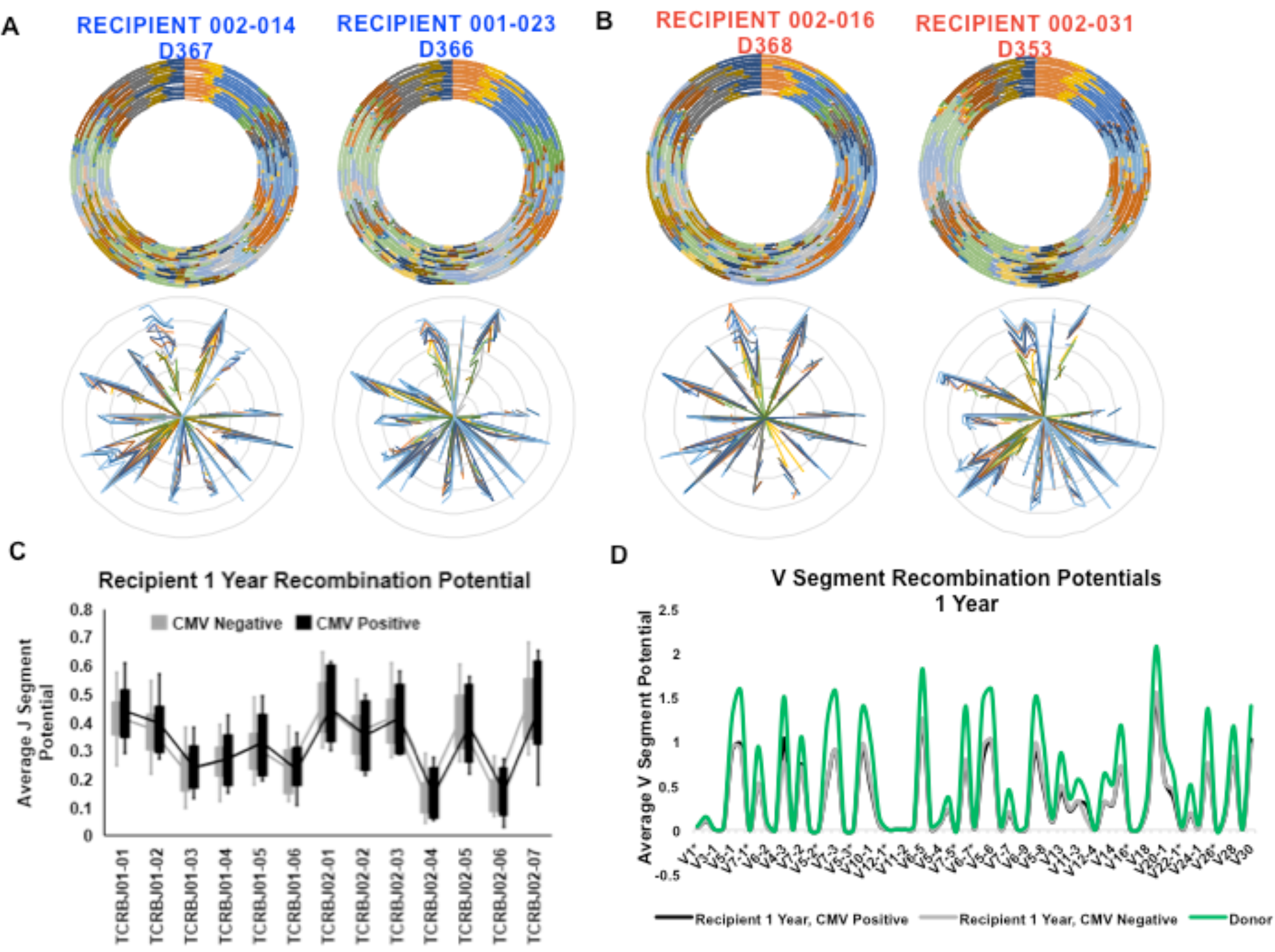
aGVHD and Recipient CMV Serostatus do not Affect Repertoire Symmetry. Additional representative RPD graphs (top) and radar plots (bottom) for patients without (A, blue) or with (B, red) aGVHD. J (C) and V (D) segment recombination potential values for patients stratified by recipient CMV serostatus (negative, n=7; positive, n=7). V segment recombination potentials for recipient at 1 month (E) or 2-3 months (F) post-BMT for patients with (n=5) or without (n=4) aGVHD with donor plotted as a control. (G) Donor V segment recombination potential values based on recipient aGVHD status.

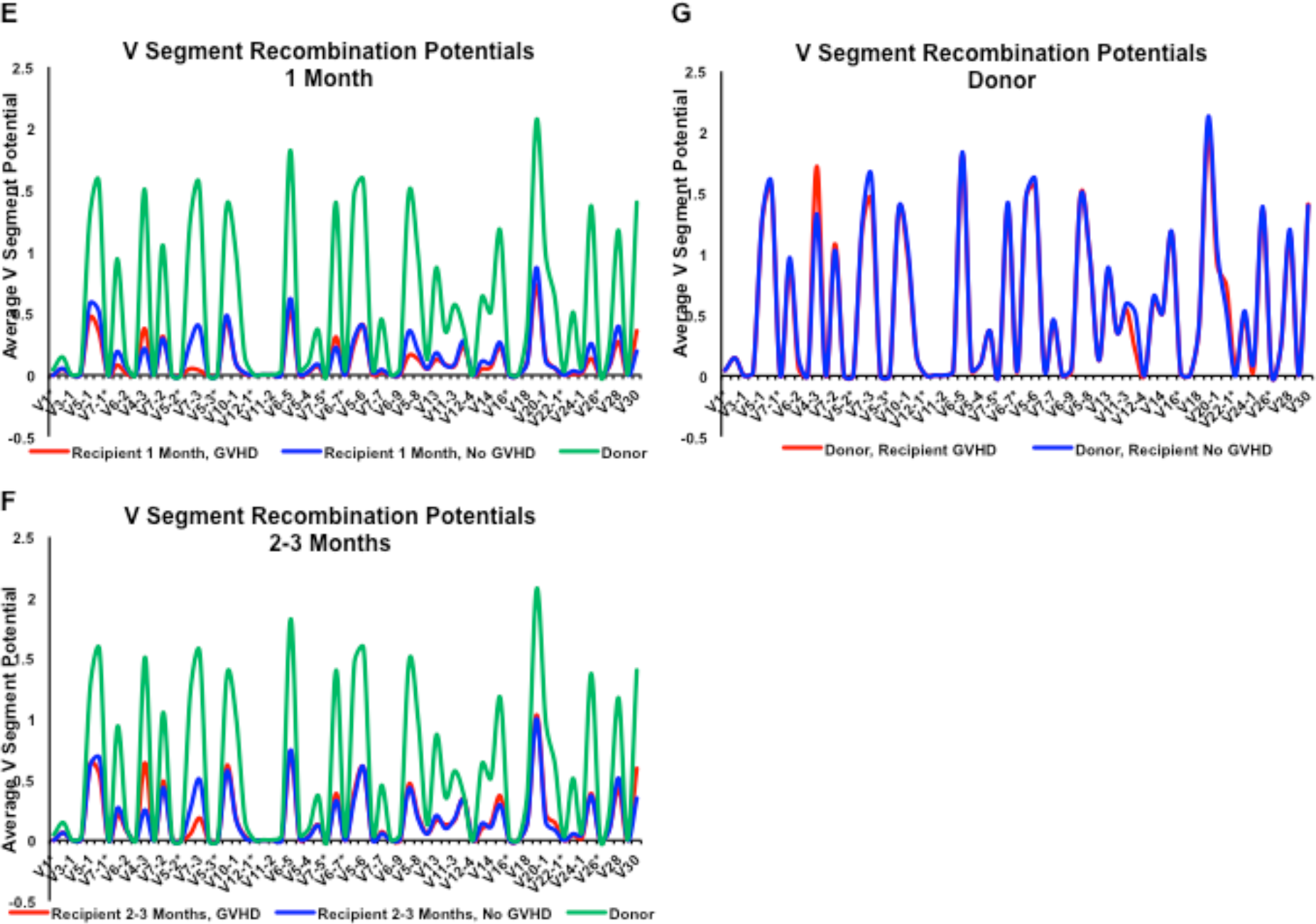

**Supplementary Figure 5:**
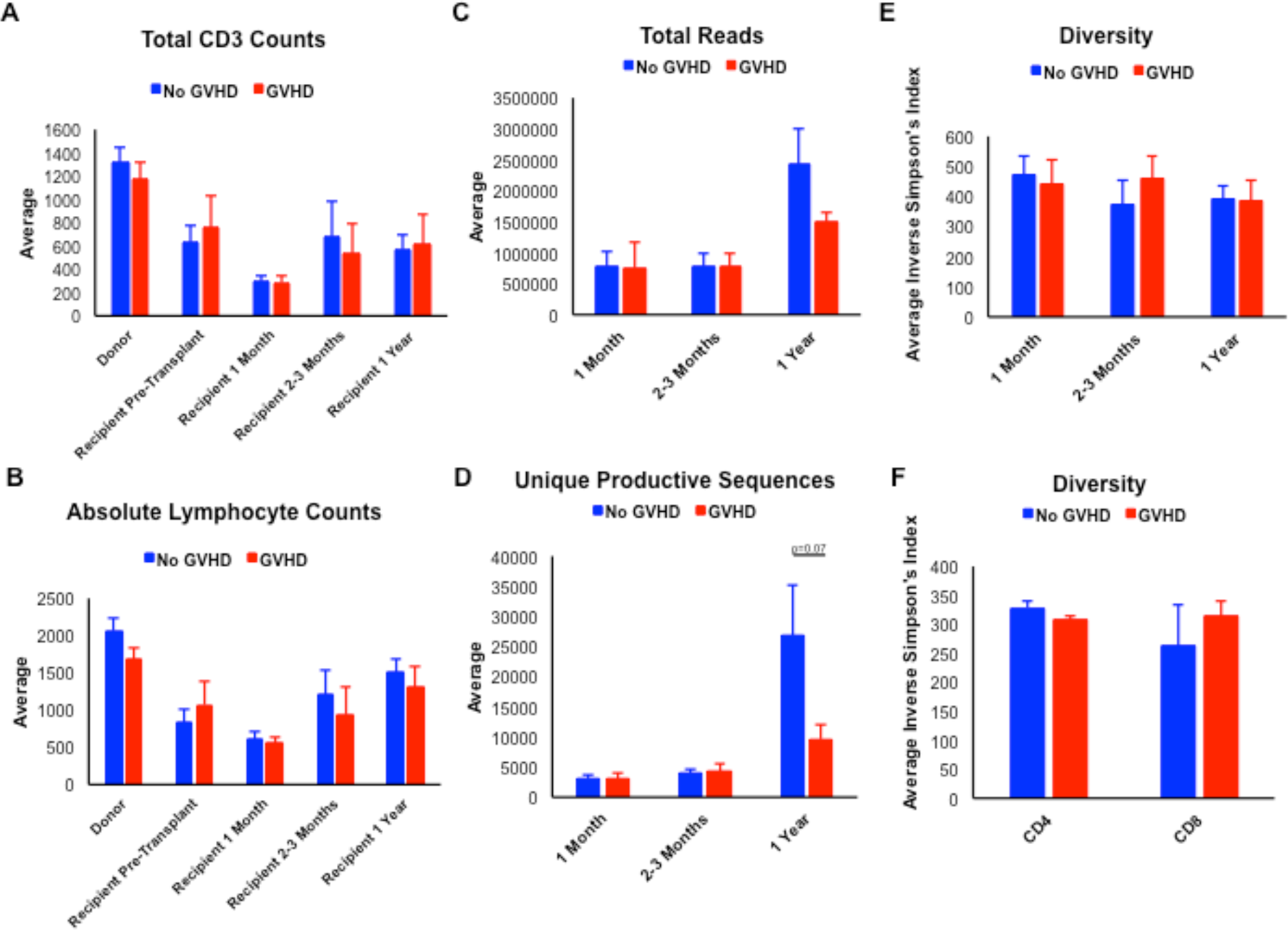
Cellular counts and sequencing parameters do not show significant differences between patients with or without aGVHD. Average total CD3 (A) or absolute lymphocyte (B) counts for BMT donors and recipients at the indicated times based on presence or absence of aGVHD. TCR sequencing results depicting total reads (C) and unique productive sequences (D) for recipients at the indicated times posttransplant based on aGVHD status (n=4-8). (E) TCR repertoire diversity values for recipients at 1 month, 2-3 months, or 1 year post-transplant for patients with (n=3) or without (n=3) aGVHD. (F) TCR repertoire diversity values for CD4^+^ or CD8^+^ T-cell subsets for patients with (n=4) or without (n=4) aGVHD.

**Supplementary Figure 6:**
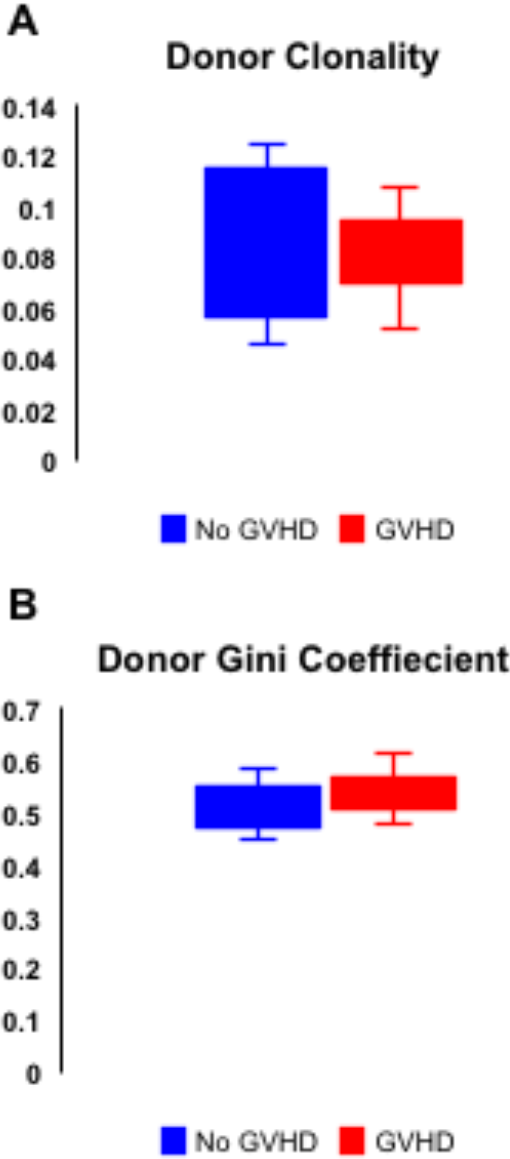
Donor Repertoires Show No Differences in Diversity based on Recipient aGVHD status. Box and whisker plot representation of donor clonality (A) and Gini coefficient (B) values based on corresponding recipient aGVHD status.

**Supplementary Figure 7:**
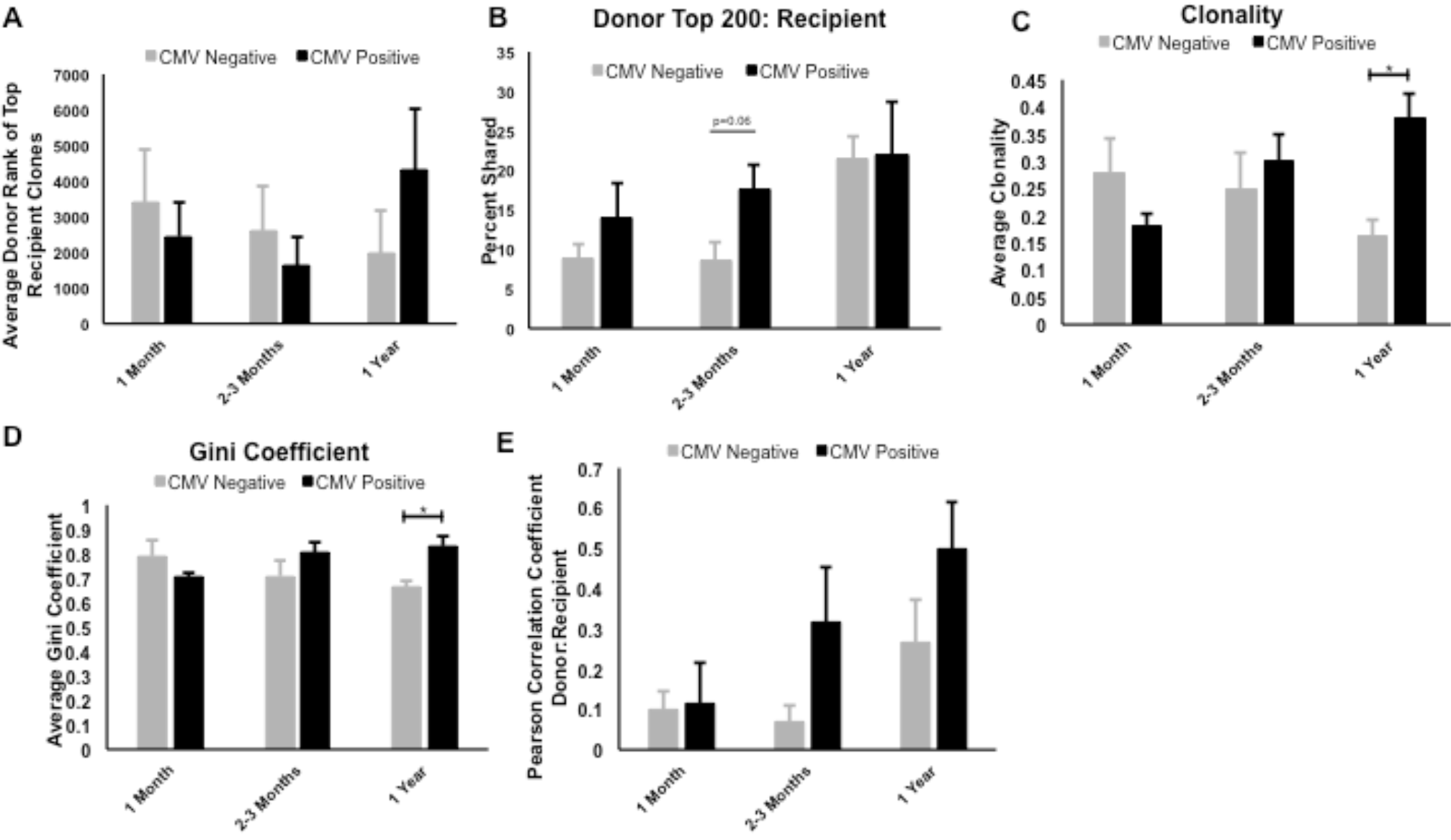
Recipient CMV Serostatus is Associated with Greater Donor:Recipient Repertoire Relatedness and Less Diversity. (A) Average donor rank of top recipient clones in patients with positive (n=5) or negative (n=4) CMV serostatus. (B) Percent of top 200 donor clones expressed in the corresponding recipient rank at the indicated time points post-transplant in patients who are seropositive (n=5) or seronegative (n=4) for CMV. p=0.06. Clonality (C) and Gini coefficient (D) values in BMT recipients at 1 month (n=4), 2-3 months (n=4-5), or 1 year (n=4) after BMT separated based on positive or negative CMV serostatus. * denotes p<0.02. (E) Quantitative analysis of the average Pearson correlation coefficient comparing donor T cell recombination events to recipients at indicated times post-transplant in patients based on CMV serostatus.

**Supplementary Table 1:** Breakdown of individual patient characteristics included in study.

| Patient | Disease | Age | Sex | Donor Type | Donor Age | Donor Sex | aGVHD<br>(Grade) | cGVHD | Patient CMV<br>Serology | Donor CMV<br>Serology | CMV<br>Reactivation | Relapse |
| --- | --- | --- | --- | --- | --- | --- | --- | --- | --- | --- | --- | --- |
| 001-001 | AML | 37 | F | MRD | 41 | M | - | - | + | - | - | - |
| 001-005 | AML | 48 | F | MRD | 56 | M | - | - | - | - | - | - |
| 001-006 | AML | 63 | M | MRD | 61 | F | II | + | - | - | - | - |
| 001-007 | AML | 60 | F | MRD | 59 | F | III/IV | - | + | - | - | - |
| 001-009 | MDS | 63 | M | MRD | 59 | F | II | - | + | - | - | + |
| 001-015 | AML | 22 | M | MRD | 17 | F | - | - | - | - | - | - |
| 001-016 | AML | 53 | F | MRD | 55 | F | II | - | - | + | - | - |
| 001-023 | ALL | 56 | F | MRD | 54 | F | - | - | - | + | - | + |
| 001-027 | AML | 60 | F | MRD | 55 | F | - | - | + | + | - | - |
| 001-035 | AML | 47 | F | MRD | 46 | M | II | - | + | - | + | - |
| 002-011 | AML | 46 | F | MRD | 38 | M | - | - | + | + | + | + |
| 002-014 | ALL | 59 | F | MRD | 57 | F | - | - | - | + | - | - |
| 002-016 | AML | 62 | F | MRD | 67 | F | II | - | + | - | + | - |
| 002-019 | AML | 33 | M | MRD | 44 | F | - | - | - | - | - | - |
| 002-031 | AML | 57 | M | MRD | 52 | M | I | - | - | - | - | - |
| 002-037 | AML | 38 | M | MRD | 35 | F | II | - | + | + | + | - |

